# Organism-wide cellular dynamics and epigenomic remodeling in mammalian aging

**DOI:** 10.1101/2025.05.12.653376

**Authors:** Ziyu Lu, Zehao Zhang, Zihan Xu, Abdulraouf Abdulraouf, Wei Zhou, Junyue Cao

**Author notes:** Correspondence (W.Z.), (J.C.). Senior author (Wei Zhou, Junyue Cao).

## Abstract

To investigate organism-wide cellular alterations and epigenomic dynamics during aging, we constructed a single-cell chromatin accessibility atlas spanning 21 mouse tissues across three age groups and both sexes. We found that around one quarter of 536 organ-specific cell types and 1,828 finer-grained subtypes exhibited significant age-related population shifts. Cellular states from broadly distributed lineages displayed synchronized dynamics with age, indicating systemic signals that coordinate these changes. Molecular analyses identified both intrinsic regulators (chromatin peaks, transcription factor activity) and extrinsic factors (cytokine programs) underlying these shifts. Moreover, ∼40% of aging-associated population dynamics were sex-dependent, with tens of thousands of peaks altered exclusively in one sex. Together, these findings present a comprehensive framework for how aging reshapes the chromatin landscape and cellular composition across diverse tissues.

## Main Text

Aging is the leading risk factor for many diseases, including cancer and neurodegenerative disorders(*1*). This association underscores the potential of therapies that target the aging process itself to delay multiple age-related diseases; however, this approach is challenging because aging is complex—each organ comprises hundreds of specialized cellular states that age differently(*2*, *3*). Systematic characterization of the cellular states most vulnerable to aging, along with their underlying genetic and epigenetic alterations, is essential for identifying therapeutic targets to guide anti-aging interventions.

Advances in single-cell transcriptomics have mapped aging-associated cellular dynamics(*4–7*), yet they largely overlook the non-coding regulation that shapes these dynamics. While single-cell ATAC-seq addresses this gap by mapping cell-type–specific chromatin changes in aging(*8–10*), most studies are limited to a few organs, leaving gaps in organism-wide inference. To overcome this limitation, here we applied EasySci-ATAC(*9*), an optimized protocol of single-cell ATAC-seq by combinatorial indexing (sci-ATAC-seq(*11*, *12*)), to define aging-associated shifts in chromatin accessibility, cell populations, and linked regulatory features—including cis-regulatory elements and transcription factor motifs—across thousands of distinct cellular states spanning different ages and both sexes.

### An organismal, single-cell chromatin accessibility atlas of aging

To map aging-associated changes of cell-type-specific chromatin landscape across the organism, we applied EasySci-ATAC(*9*) to profile 552 samples from 32 mice spanning three ages (1, 5, and 21 months) with 8–12 sex-balanced replicates per group (except ovary/uterus) (**Table S1**). A total of 21 tissue types across major systems were analyzed, except for the brain, which had been analyzed in our previous study(*9*). To support a companion study of newborn cells, we labeled mice with 5-Ethynyl-2-deoxyuridine (EdU) before tissue collection, similar to our previous work(*10*). This allows for parallel profiling of DAPI singlets (“all” cells; this study) and EdU+ nuclei (“newborn” cells, companion study).

We extracted nuclei from each tissue, followed by sorting, barcoding, and sequencing, yielding ∼95 billion paired-end reads. After filtering out low-quality cells and doublets (**Fig. S1A-C**), we obtained 10,956,311 single-cell profiles, including 6,839,086 DAPI singlets and 4,117,225 EdU+ cells (**Fig. 1B and Fig. S1D**). With a low duplication rate (20.1%), we detected a median of 3,031 unique fragments per cell; 29.8% of reads mapped to promoter regions (within ±1 kb of the transcription start site, TSS), consistent with prior scATAC studies(*9*, *13*) (**Fig. S1E-F**). Pseudobulk profiles clustered by tissue type rather than mouse, indicating low batch effects (**Fig. S1G**).

**Figure 1.**
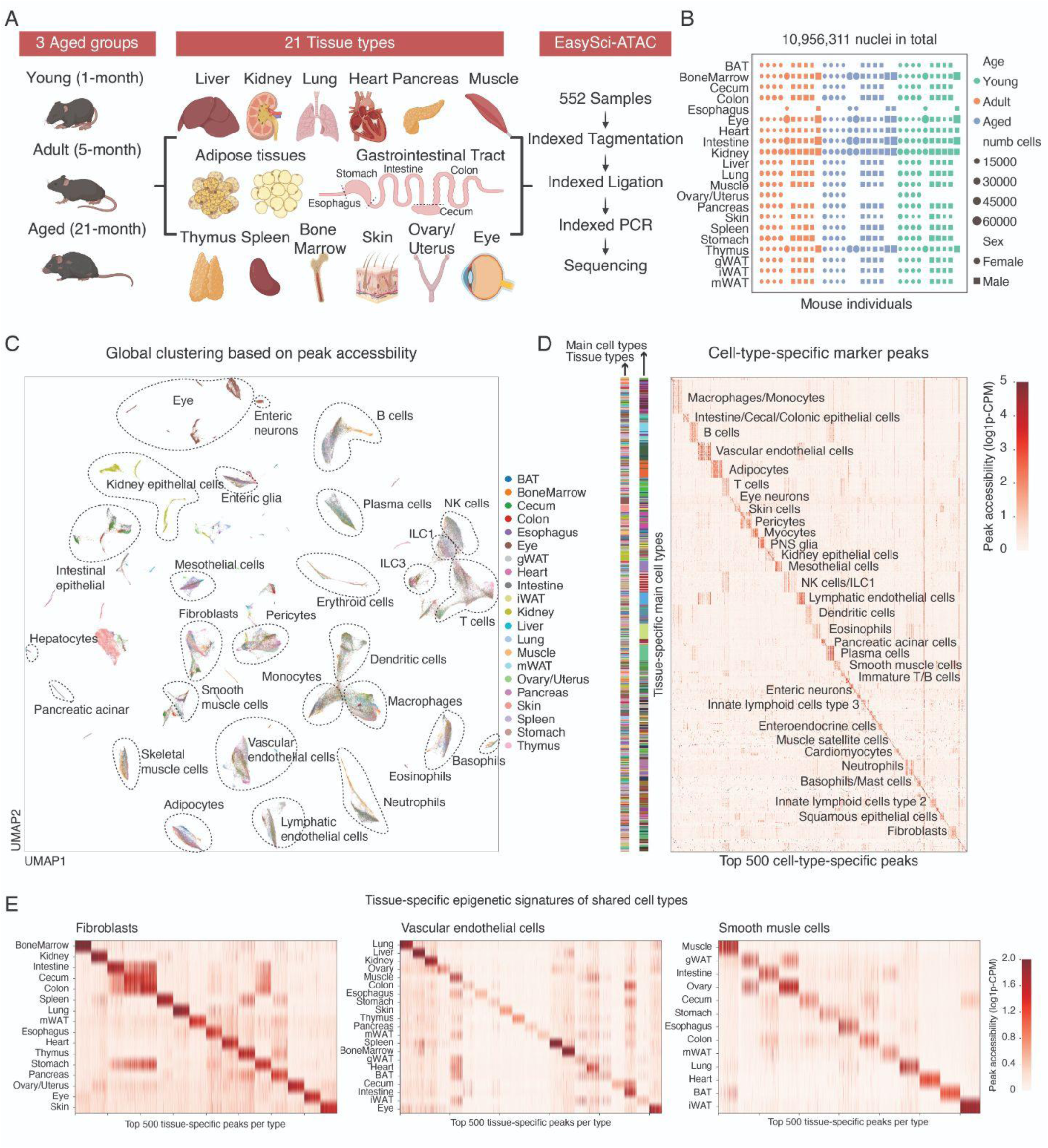
An organismal, single-cell chromatin accessibility atlas of aging. **(A)** Experimental scheme to construct an organismal cell atlas of chromatin accessibility across different ages and both sexes. **(B)** Dot plot showing the total number of cells obtained from each mouse individual. **(C)** UMAP visualization of the entire dataset subsampled to a maximum of 800 cells per main cell type per tissue, colored by the tissue type. Dimension reduction was performed using the peak-count matrix. The same cell types from multiple tissues that clustered together were circled. **(D)** Heatmap showing the aggregated accessibility of peaks specific to each main cell type, quantified by counts per million. Cell-type-specific peaks were identified using an entropy-based strategy (*18*). The top 500 peaks were selected for each main cell type, ranked by specificity score. **(E)** Heatmap showing the aggregated accessibility of peaks specific to each tissue for broadly distributed cell types, identified similarly to D.

We identified different cell types per tissue using SnapATAC2(*14*) for dimension reduction and Leiden clustering. This was followed by manual annotation based on the accessibility of cell markers (**Table S2, Fig. S2**), revealing 144 unique cell types (e.g., T cells, B cells), or 536 organ-level cell types (e.g., lung B cells, spleen B cells) (**Fig. S3A**). While most unique cell types were organ-restricted (n = 102), 18 cell types were shared across >10 tissues, such as immune cells (*e.g.,* T cells, B cells), stromal cells (*e.g.,* fibroblasts, smooth muscle cells), and endothelial cells (*e.g.,* vascular and lymphatic endothelial cells). We further confirmed the reproducibility of our annotations through a leave-one-out cross-validation strategy (*15*) (**Fig. S3B-D**) and validated the cell type identities through integration analysis with scRNA-seq atlases (*7*) (**Fig. S4**).

To define a universally accessible map, we merged the accessible peaks called from each cell type within the tissue, yielding a total of 1,341,077 open chromatin regions (±250 bp from summits). The genomic distribution of these peaks was largely consistent with previous sn-ATAC-seq profiling (*16*) (**Fig. S5A**): they were predominantly located within introns (n=648,244; 48.3%) and intergenic regions (n=404,499; 30.1%), and to a lesser extent at TSS-promoter regions (n=34,487; 2.57%). These regions occupy 12.3% of the genome, intersecting with 89.1% of CREs (cis-regulatory elements) identified by the ENCODE consortium (*17*) and 88.9% of CREs from Cusanovich et al (*12*)(**Fig. S5B**). Using this peak set, cells are clustered by lineage across tissues, further supporting our annotations (**Fig. 1C**).

We next investigated the cell-type-specific usage of CREs. We first performed NMF decomposition (*16*) on the aggregated cell type by peak matrix (**Fig. S5C**). The majority of CRE modules were restricted to a specific subset of cell types (**Fig. S5D**). We then followed up with an entropy-based method (*18*) to identify CREs that are specific at different hierarchical levels. At the whole-organism level, shared cell types across tissues display conserved epigenomic signatures (**Fig. 1D**), while tissue-specific signatures became evident when examined at higher resolution (**Fig. 1E**). We next applied chromVar (*19*) to pinpoint TF motifs specific to each cell type, such as the enrichment of *Runx3* in cytotoxic T cells and NK cells (*20*), *Sfpi1* in general myeloid cells (*21*), *Irf4* in plasma cells (*22*), and *Ebf1* in B cells and pericytes (*23*, *24*) (**Fig. S5E**). The enrichment of these motifs was further validated by gene activity and cross-validated across different organs (**Fig. S5F**).

Next, we performed linkage analysis using matched cell types from published RNA-seq (*7*) to identify potential target genes of CREs (**Fig. S6A–B**). We identified a union of ∼2.1 million positively correlated gene–peak pairs across age groups. These linkages were highly reproducible (72% detected in at least two age groups; **Fig. S6C**) and displayed coordinated chromatin accessibility with expression of their putative target genes (**Fig. S6D**).

Lastly, we sought to use cell-type-specific accessible peaks to interpret genetic variants associated with complex traits and diseases. Specifically, we lifted over human GWAS variants to mice and then applied linkage disequilibrium score regression (LDSC) to assess GWAS SNP enrichment within cell-type-specific accessible peaks(*25*) (**Methods**). Enrichment matched known biology: multiple sclerosis in B/T/plasma cells (*26*, *27*), LDL in hepatocytes(*28*), height in tenocytes, type 2 diabetes in pancreatic β cells (*29*), and hypertension in kidney juxtaglomerular cells(*30*) (**Fig. S5G**).

### Age- and sex- dependent cell population change at the main cluster level

We observed widely varied fractions of different cell types across tissues (**Fig. 2A**): 43 organ-restricted types comprised <1% of their tissues, such as *Ascl1*+ lung neuroendocrine cells (0.039%) and *Gja8*+ lens epithelium (0.051%), highlighting the importance of throughput for identifying these rare yet critical populations. We then assess sex dimorphism across cell types. While most cell types exhibited similar population sizes between sexes (**Fig. S7A**), we identified a subset of cell types displaying strong sex-specific differences in chromatin landscapes. Specifically, we trained k-nearest neighbor classifiers for each cell type to distinguish female vs. male cells from scATAC-seq embeddings, and then used the area under the curve (AUC) metric to identify cell types with differences between genders, such as hepatocytes, proximal tubule cells, type IIB myonuclei, and multiple gonadal WAT types (**Fig. 2B**), mirroring previous reports (*7*, *31*, *32*). For example, proximal tubule (PT) cells in the kidney, including the general states spanning spatially connected segments (PTS1, PTS2, and PTS3; marked by *Slc34a1*) and a distinct state of segment 3 (PTS3T2, marked by *Slc22a7*) (*33*) was separated into clusters determined by sexes with unique marker gene accessibility (**Fig. 2C and Fig. S7B**). Sex differences were widespread on autosomes and highly cell-type-specific (**Fig. 2C, right**). TF Motif analysis of sex-biased regions, validated with matched scRNA-seq (**Fig. S7C**), revealed elevated HOX/KLF motifs in male hepatocytes (**Fig. S7D–E**) and androgen-response elements in male PT cells (**Fig. S7F–G**), indicating lineage-specific routes to sex-biased regulation.

**Figure 2.**
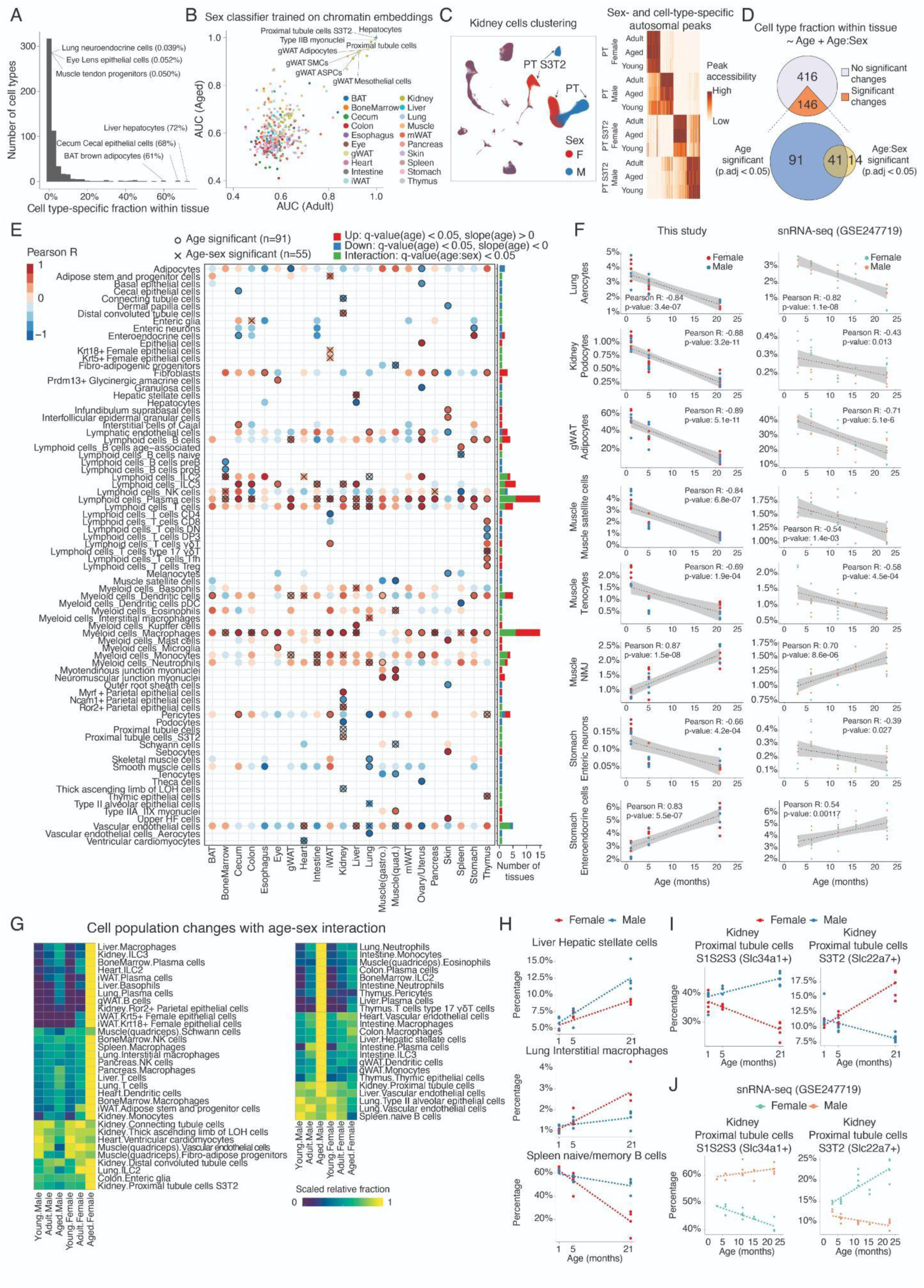
Age- and sex-dependent cell population changes at the main cell type level. **(A)** Histogram showing the distribution of cell-type-specific fractions within tissue. Only cells from adult mice (5-month-old) were used in this quantification. **(B)** K-means classifiers were trained to distinguish male and female cells for each main cell type using spectral embedding calculated by SnapATAC2 (*14*). Scatterplot comparing the area under the curve (AUC) values of models between adult and aged cells. Cell types with high sex dimorphism in both age groups are labeled. ASPC, adipose stem and progenitor cells. **(C)** Left: UMAP plots of all kidney cells, colored by sex (right), highlighting distinct chromatin states between sexes of proximal tubule cells (PT). Right: Heatmap displaying accessibility of autosomal peaks with sex-and cell-type-specific patterns, conserved across ages. **(D)** Venn diagram showing the number of main cell types whose relative proportions in corresponding tissues are significantly associated with age or age-sex interactions. **(E)** Left: Dotplot showing all significantly altered main cell types from (D) across tissues, colored by Pearson correlation between their relative proportion and age. Cell types with significant age or age-sex interactions are marked. Right: Barplot summarizing the number of tissues with significant changes for each cell type, colored by direction (up/down) or interaction. **(F)** Scatterplot illustrating examples of main cell types with significant age-related proportional changes (left), alongside validation using published snRNA-seq data (*7*) for the same cell type and tissue (right). **(G)** Heatmap showing relative proportions of main cell types with significant age-sex interactions, including female-biased (left) and male-biased (right) cell types. Subtype fractions were first calculated for each animal (normalized to total cells recovered per tissue), and averaged in each tissue/sex/age group, and then scaled the values to [0,1]. Significant age-sex interactions were defined as q-value (age:sex) < 0.05 and coefficient of determination (R²) > 0.4. **(H)** Scatterplot highlighting cell types that change in the same direction with aging but exhibit sex-dependent magnitude. Normalization was performed at the level of individual biological replicates (i.e., within each mouse). **(I-J)** Scatterplots of two kidney proximal tubule cell types with opposite sex-specific proportional trends (I) and snRNA-seq validation results (J).

We next investigated the effects of aging on cell population dynamics by fitting linear models with age and age×sex terms per organ (**Methods**). This revealed 146 organ-specific cell types significantly altered with age (FDR ≤ 0.05, R² > 0.4), 55 of which showed significant sex interactions (**Fig. 2D–E; Table S3**). While all organs exhibited changes, ovary/uterus, liver, and thymus showed the highest fractions of affected cell types (**Fig. S8A**). 62 organ-specific cell types expanded with age regardless of sex (**Fig. 2E, Fig. S8B top**), 68% of which were from immune lineages (**Fig. S8C**). These include broadly expanded cell types (*e.g.,* plasma cells and macrophages) as well as tissue-restricted expansions such as Kupffer cells (liver) and γδ T cells (iWAT) (**Fig. S8D**), consistent with previous studies(*34–38*). We also detected expansion of non-immune cell types, such as increased stomach enteroendocrine cells (**Fig 2F left)**, potentially contributing to dysregulated digestive processes in aged individuals (*39*).

The 29 aging-depleted cell types were largely tissue-specific (**Fig. 2E, Fig. S8B bottom**). This includes immune progenitors such as bone-marrow pro-/pre-B cells and thymic T cell precursors, consistent with the decreased lymphopoiesis(*40*) and thymopoiesis (*41*). Meanwhile, we detected multiple non-immune cell types that are significantly depleted especially in the ovary/uterus, skin, and lung (**Fig. S9A**): aged ovary/uterus showed reduced population of granulosa and theca cells, basal epithelial cells and vascular endothelial cells (**Fig. S9B**), consistent with follicle depletion in reproductive aging(*42*, *43*). Aged skin exhibited reductions in melanocytes, dermal papilla cells, outer root sheath cells, and lymphatic endothelial cells (**Fig. S9B**), the latter consistent with reduced lymphatic density in aging skin(*44*). The aged lung showed declines in vascular-associated stromal cells (pericytes, smooth muscle cells) and aerocytes(*45*) (**Fig. S9B**), suggesting compromised blood flow regulation with age(*46*).

Other depleted functional cell types included kidney podocytes and *Ncam1*+ parietal epithelial cells (a group of potential podocyte progenitors(*47*)) (**Fig. 2E-F, Fig. S9B)**, indicating impaired function and regenerative capacity in the aged kidney. Additionally, we observed depletion of diverse cell types, including liver hepatocytes (*48*), muscle tenocytes(*49*), muscle satellite cells (*50*), stomach enteric neurons, and adipocytes from multiple organs (*e.g.,* lung, gWAT, and ovary/uterus) (**Fig. 2E-F and S9B)**. Some cell types exhibited over 50% reductions by 5 months (vs. 1 month), such as satellite cells, tenocytes, and pro-/pre-B cells, highlighting an early onset of aging impacts on the population of these cell types.

We further analyzed 55 cell types displaying significant age-sex interactions. Most sex-specific cell types (35 out of 55) are immune cells: 15 cell types expanded preferentially in aging males, including neutrophils (lung, intestine) and eosinophils (muscle), consistent with human studies (*51*, *52*), whereas aged females showed increased basophils (liver) and NK cells (pancreas, bone marrow) (**Fig. 2G)**. Additionally, some cell types exhibited similar trends in both sexes but differed in the extent of change in different organs (*e.g.,* male-biased expansion of bone-marrow ILC2s and hepatic stellate cells; female-biased lung interstitial macrophage expansion and spleen naïve/memory B-cell decline, **Fig. 2G-H**). Interestingly, certain cell types showed opposite trends between the sexes. For instance, the aforementioned specialized proximal tubule (PT) cell type (PT S3T2, characterized by *Slc22a7*) increased in females but decreased in males, whereas the other PT cells (PT S1S2S3, marked by *Slc34a1*) exhibited the opposite pattern (**Fig. 2I)**.

Lastly, we validated the age-related population shifts identified in our scATAC-seq data using the published PanSci scRNA-seq dataset (*7*) (**Fig. S10A**). Among 68 cell types with scRNA-seq counterparts, 71.4% of aging-expanded populations (**Fig. S10B**) and 89.5% of aging-depleted populations (**Fig. S10C**) exhibited concordant trends. Notable expansions include lymphoid cells (T cells, B cells, and plasma cells) across tissues (**Fig. S10B**), NMJs in muscle, and enteroendocrine cells in the stomach (**Fig. 2F right**). Consistent depletions were also observed, such as lung aerocytes, muscle satellite cells and tenocytes, and kidney podocytes (**Fig. 2F right**). In addition, we confirmed sex-biased dynamics in 12 cell types, such as female-biased expansion of B cells in gWAT and T cells in the liver, as well as opposing trajectories between two kidney PT cell types (**Fig. 2J**). Together, these cross-modality validations demonstrate that our scATAC-seq findings capture conserved and reproducible aging trajectories.

### Age- and sex- dependent cell population change at the subtype level

We further explored cell population dynamics at the subtype level. First, we re-clustered each main cell type (with >200 cells) using SnapATAC2 (*14*), and manually curated cellular subtypes based on cluster-specific gene accessibility, revealing substantial heterogeneity within main cell types. For example, eye amacrine cells were clustered into 34 subtypes with distinct chromatin landscapes and TF activity (**Fig. 3A, Fig. S11A-B**), aligning with scRNA-seq studies(*53*). Across the organism, we identified 1,828 subtypes from 386 main types, with a median of 1,610 cells per subtype and four subtypes per main cell type (**Fig. S11C**). Most subtypes (93.4%) were consistently detected across more than ten samples (**Fig. S11C**). This includes rare subtypes underexplored in previous scATAC-seq studies, such as small-intestine enteroendocrine L cells (*Gcg+*, 0.092% of the intestine cell population), K cells (Gip, 0.070%), D cells (*Sst+*, 0.028%), enterochromaffin (*Tph1+*, 0.084%), and *Neurog3*+ progenitors (0.081%) (**Fig. S11D-E**).

**Figure 3.**
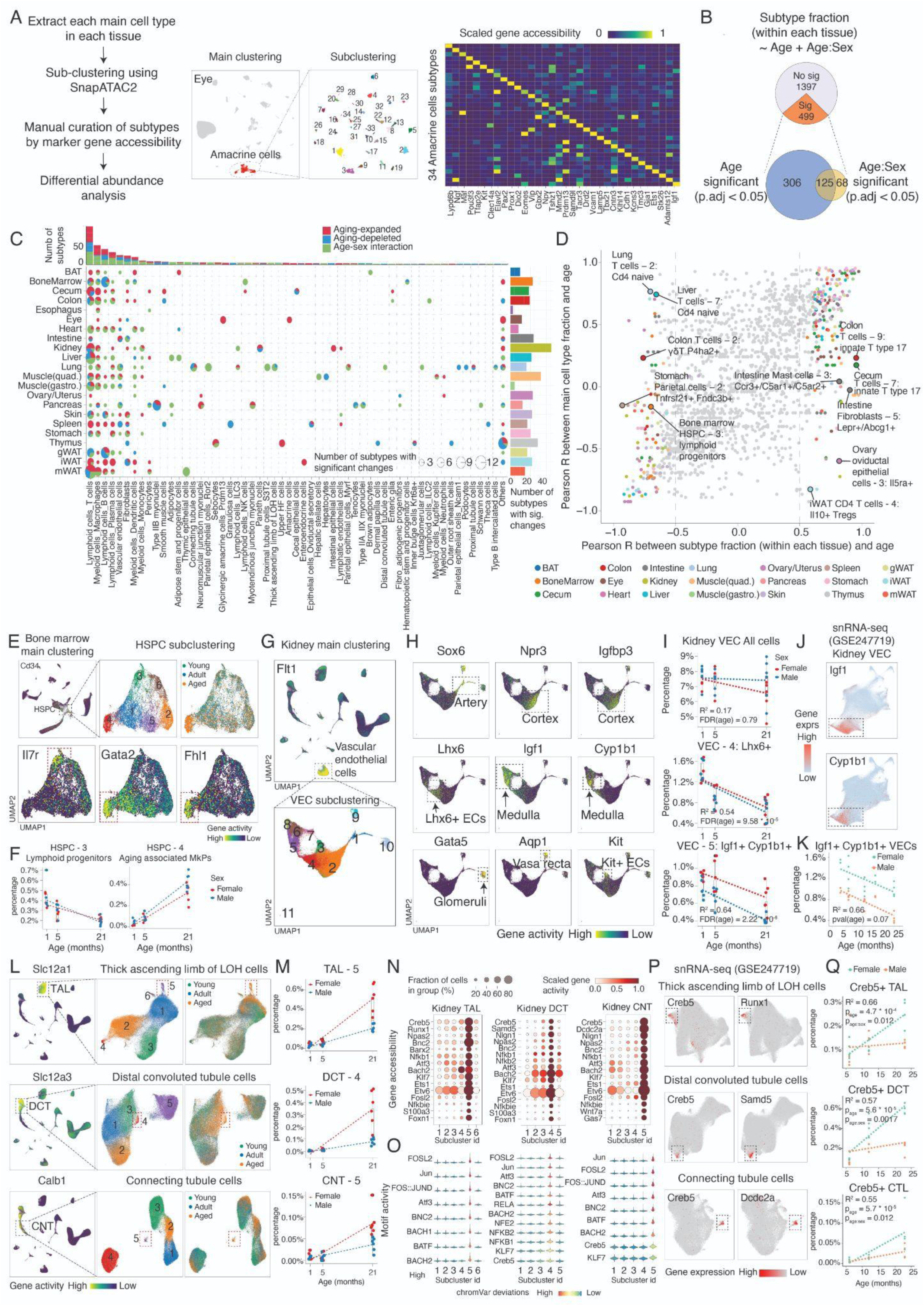
Age- and sex-dependent cell population change at the subtype level. **(A)** Left: schematic of the sub-clustering workflow. Middle: example UMAP plots showing sub-clustering analysis of eye amacrine cells, colored by subcluster ID. Right: heatmap displaying marker gene accessibility for amacrine cell subtypes. Marker genes were determined using a t-test with overestimation of variances in Scanpy(*75*), with a significance cutoff of q < 0.05. **(B)** Venn diagram showing the number of cell subtypes with significant changes associated with age or age-sex interactions. **(C)** Top: barplot showing the number of aging-associated subtypes for each main cell type. Center: dotplot summarizing the number of aging-associated subtypes across main cell types and tissues. Right: barplot showing the number of aging-associated subtypes per tissue. **(D)** Scatterplot comparing aging-associated cellular fraction changes between aging-associated subtypes and their corresponding main cell types. Subtypes with distinct trends compared to their parent main cell types are labeled. Subtype fractions were calculated relative to the whole tissue. **(E)** UMAP plots showing sub-clustering of bone marrow HSPCs to identify different subtypes, colored by subcluster ID (top middle), age group (top right), and marker gene accessibilities for lymphoid progenitor cells (bottom left) and age-associated megakaryocyte progenitors (bottom right). **(F)** Scatter plot showing cell proportion changes for lymphoid progenitor cells (HSPC-3) and age-associated megakaryocyte progenitors (HSPC-4), with a linear regression fit. Each dot represents one animal. **(G)** UMAP plot showing sub-clustering results of kidney vascular endothelial cells to identify different subtypes, colored by subcluster ID (bottom). **(H)** UMAP plots of kidney vascular endothelial cells, colored by the accessibility of genes that mark vascular cells across different spatial locations. **(I)** Scatter plots and boxplots showing the fractions of all kidney endothelial cells (top), vein endothelial cells (*Lhx6*+ VEC-4, center), and medulla endothelial cells (*Igf1*+ *Cyp1b1*+ VEC-5, bottom) across three age groups. **(J)** UMAP plots of vascular endothelial cells from a published snRNA-seq study (*7*), colored by the expression of genes marking medulla endothelial cells (as in **H**). **(K)** Scatterplot showing the aging-associated decline in the kidney medulla endothelial cells identified from a snRNA-seq atlas of aging(*7*). **(L)** UMAP plots showing all kidney cells (Left) and sub-clustering analysis (middle and right) for thick ascending limb cells (TAL, top), distal convoluted tubule cells (DCT, center), and connecting tubule cells (CNT, bottom), colored by main cell type specific markers (left), subtype ID (middle) and age group (right). Squares indicate the aging-associated cell subtypes. **(M)** Scatter plots showing the relative fractions of three reactive subtypes identified in (**K**) across ages, with separate linear regression fits for each sex. **(N)** Dotplots showing the gene marker accessibility marking the reactive states (TAL-5, DCT-4 and CNT-5) in each renal epithelial cell type. Marker genes were determined using a t-test with overestimation of variances in Scanpy (*75*), with a significance cutoff of q < 0.05. **(O)** Violinplots showing the motif accessibilities of transcription factors marking the reactive states in each renal epithelial cell type. **(P)** UMAP plots showing sub-clustering results for TAL, DCT, and CNT cells from published snRNA-seq data(*7*), colored by the expression of genes that mark reactive states (as in **N**). **(Q)** Scatterplots showing the relative fractions of reactive subtypes identified in **(P)** across aging, with separate linear regression fits for each sex.

Using the same analytical pipeline as for main cell types, we identified 499 subtypes (out of 228 organ-specific main cell types) with significant age-associated population dynamics (FDR of 0.05, R^2^ > 0.4) (**Fig. 3B, Table S4**), including 193 subtypes with sex-specific aging differences (FDR of 0.05) (**Fig. 3C**). Tissues harbored a median of 24 altered subtypes, ranging from 3(esophagus) to 51 (kidney). While most subtypes exhibit similar dynamics with their parent main cell types (Pearson r: 0.72, p-value < 2.2 * 10^-26^), we observed many exceptions: specific aging-associated subtypes such as *Lepr1+ Abcg1+* fibroblasts and *Ccr3+ C5ar1+ C5ar2+* mast cells expanded in the aged intestine, while their main cell types remained relatively stable (**Fig. S11F-G**). In the liver and lung, the overall main T cell population increased in aged tissues, yet subtypes corresponding to naive T cells were depleted (**Fig. S11F-G**). We also observed opposing aging-associated dynamics of subtypes from the same cell type. For example, in bone marrow HSPCs, lymphoid progenitors (HSPC-3, *Il7r*+) declined while megakaryocyte-biased HSPCs (HSPC-4, *Gata2/Fhl1/Hoxb5*+) increased (**Fig. 3E-F)**, consistent with a recent study(*54*).

Analysis of these aging-associated subtypes revealed region-specific vulnerability of the same cell type. For example, renal vascular endothelial cells (ECs) segregated into subtypes representing ECs from different anatomical locations(*55*), including the artery (*Sox6+*), cortex (*Npr3+, Igfbp3+*), vein (*Lhx6+*), medulla (*Igf1+, Cyp1b1+*), vasa recta (*Aqp1+*), and glomeruli (*Gata5+*) (**Fig. 3G-H**). While the total abundance of ECs remains relatively constant, we observed a significant depletion of two EC subtypes—*Lhx6+* vein ECs(*56*, *57*) and *Igf1+, Cyp1b1+* medulla ECs(*55*) (**Fig. 3I**). Re-analysis of a recent snRNA-seq atlas(*7*) further confirmed the decline of *Igf1+, Cyp1b1+* medulla ECs in both sexes (with a higher female baseline) (**Fig. 3J-K)**.

Meanwhile, we found shared “reactive” features across different aging-expanded subtypes. For example, sub-clustering of three specialized renal epithelial cell types—the thick ascending limb of the Loop of Henle (TAL), distal convoluted tubule (DCT), and connecting tubule (CNT)—revealed three subtypes (*i.e.,* TAL-5, DCT-4, and CNT-5) that exhibit significant expansion in aging, particularly in females (**Fig. 3L-M)**. All three subtypes exhibited shared activation of TFs linked to inflammation and stress responses (*e.g., Creb5, Bnc2, Runx1, Bach2, Fosl2,* and *Klf7*), supported by gene-body accessibility and motif activity (**Fig. 3N–O**). Re-analysis of a scRNA atlas(*7*) validated both the state and its female-biased expansion (**Fig. 3P–Q**), implicating inflammatory signaling in the aged female kidney.

Lastly, to systematically evaluate subtypes identified from the current study that reflect biologically meaningful entities, we used scBridge(*58*) to transfer subtype annotations from scATAC-seq to scRNA-seq, and examined the results through two orthogonal criteria: molecular similarity and population dynamics consistency (**Fig. S12A**). Reassuringly, matched subtypes showed consistently higher RNA–ATAC correlations than unmatched pairs (**Fig. S12B**), and aging-associated changes were largely concordant across both modalities (**Fig. S12C**). As illustrative examples, we highlight kidney juxtaglomerular cells, stomach enteroendocrine cells, kidney vascular endothelial cells, and intestinal enteric neurons (**Fig. S12D–G**), where subtypes not only showed strong molecular similarity but also displayed parallel age-related compositional trajectories across two datasets.

### Aging-associated population dynamics of broadly distributed cell subtypes

We next examined whether broadly distributed lineages (immune and stromal) displayed coordinated aging dynamics across organs. Starting from immune cells, we applied clustering analysis to T cells (1,264,322), B cells (1,079,641), plasma cells (157,545), NK/ILC1/2/3 (87,550), monocytes/macrophages (581,877), and dendritic cells (306,562), revealing 67 subtypes defined by distinct accessibility at canonical markers (**Fig. 4A-B, Fig. S13A**). Many subtypes were highly tissue specific (**Fig. 4C and Fig. S14**): thymic DN/DP T cells; intestinal Gzma+ γδ T cells; bone-marrow B progenitors; gut-enriched IgA+ plasma cells. For macrophages (MAC), *Cxcl16*+ *Tlr12*+ macrophages (MAC-6/7) were constrained to the small and large intestine, whereas *Spic*+ macrophages were unique to bone marrow (MAC-9) and spleen (MAC-10) (**Fig. 4C**), consistent with the role of *Spic* in iron-recycling macrophage development (*59*). Certain dendritic cells also demonstrated tissue-specific characteristics, such as *Apol7c*+ type I DCs (DC-3), *Il22ra2*+, and *C1qtnf3*+ type II DCs (DC-9/10), which were exclusively identified in the gut (**Fig. 4C**).

**Figure 4.**
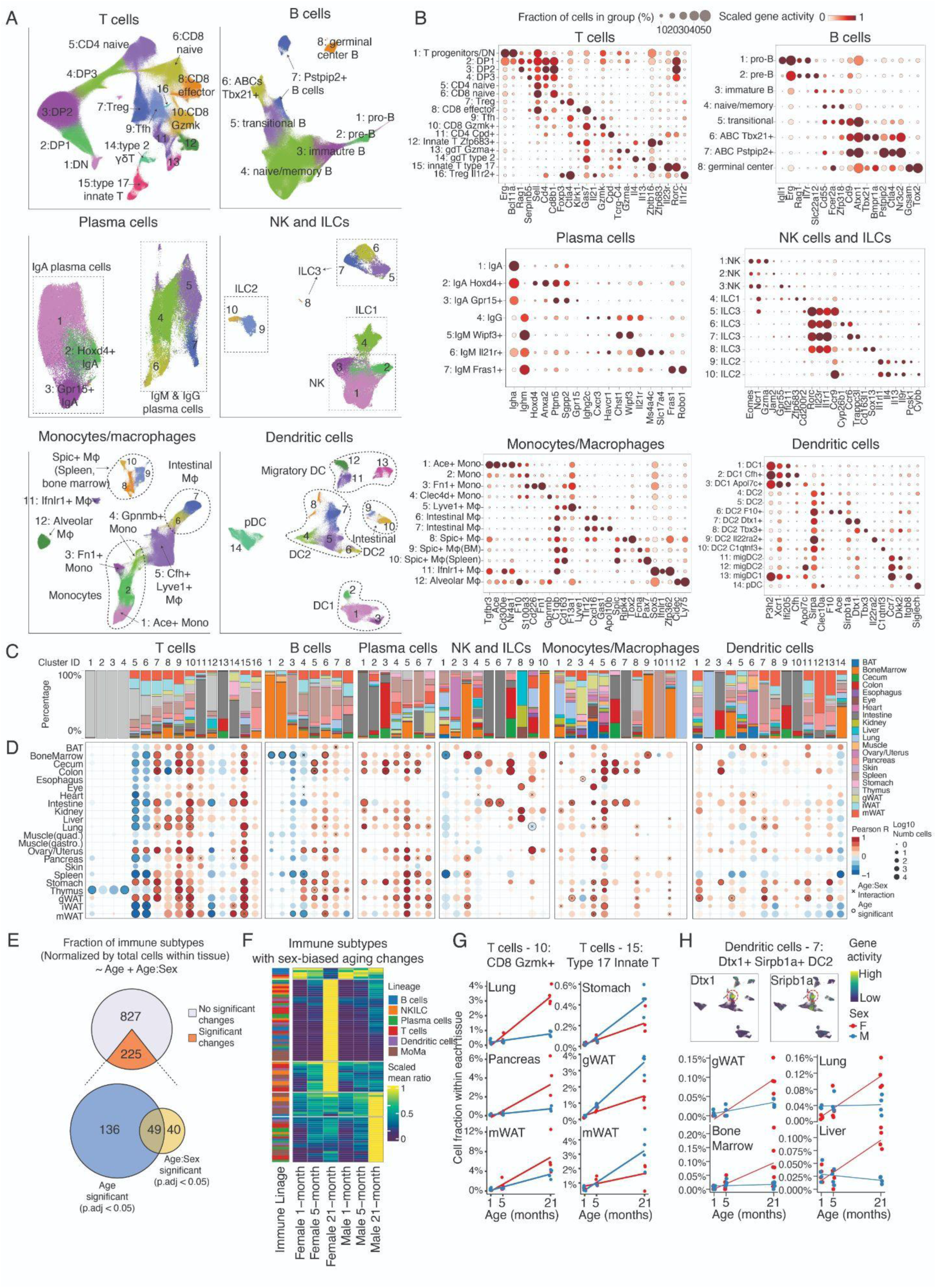
Aging-associated dynamics of broadly distributed immune subtypes. **(A)** UMAP plots showing combined clustering results for T cells, B cells, plasma cells, innate lymphoid cells, macrophages, and dendritic cells across tissues, colored by subtype ID. **(B)** Dotplots showing accessibility of genes used for annotating immune cell subtypes. **(C)** Stacked bar plots showing the tissue composition for each immune subtype. **(D)** Dotplots showing the proportional changes of each immune subtype in each tissue across aging, colored by Pearson correlation between their relative proportion (within each tissue) and age. Subtypes with significant age or age-sex interactions are marked. **(E)** Venn diagram showing the number of cell subtypes with significant changes associated with age or age-sex interactions. **(F)** Heatmap showing relative proportions of immune cell subtypes with significant age-sex interactions, including female-biased (left) and male-biased (right) cell types. Subtype fractions were first calculated for each animal (normalized to total cells recovered per tissue), and averaged in each tissue/sex/age group, and then scaled the values to [0,1]. Significant age-sex interactions were defined as q-value (age:sex) < 0.05 and coefficient of determination (R²) > 0.4. **(G)** Scatterplots showing the female-biased expansion of *Gzmk+* CD8 T cells (left) and male-biased expansion of *Il23r+ Zbtb16*+ type 17 innate T cells (right) along aging. Each dot represents one animal, with linear regression lines added for each sex. Subtype fractions were normalized by total cells within each tissue for each sample. **(H)** UMAP plots of *Dtx1*+ *Sirbp1a*+ type 2 dendritic cells colored by accessibility of marker genes. Scatter plot showing their female-biased expansion along aging, with separate linear regression fits for each sex.

Differential proportion analyses (**Fig. 4D-E, Table S5**) showed broad depletion of naïve CD4⁺ (in 10 tissues) and naïve CD8⁺ T cells (in 5 tissues), immature T cells in thymus, and pro-/pre-B cells in bone marrow, with corresponding declines of naïve/memory B cells in spleen and mWAT (**Fig. 4D**). Additional losses included *Zfp683⁺* Trm cells in intestine/iWAT/mWAT and *Gzma⁺* resting NK cells in spleen/bone marrow, coinciding with expansion of *Gpr55⁺ Ifi211⁺* proinflammatory NK cells in the same organs (**Fig. 4D**). In contrast, 42/67 immune subtypes expanded with age in at least one tissue, spanning T-cell (*e.g., Zbtb16⁺ Il23r⁺* type-17; *Gzmk⁺* CD8; *Il21⁺* Tfh; *Foxp3⁺* Tregs), B-cell (*Tbx21⁺* ABCs), plasma (*Chst1⁺ Wipf3⁺* IgM), monocyte/macrophage (*Clec4d⁺* monocytes; *F13a1⁺ Lyve1⁺* macrophages), and multiple DC subtypes (**Fig. 4D**). Some expansions were gut-specific (*e.g., Gpr15⁺ Stc2⁺* IgA plasma; *Tlr12⁺* MAC-7), and others showed organ-dependent polarity (*e.g., Klrk1⁺ Gas7⁺* CD8 effectors increased in liver/lung but declined in colon/cecum), reflecting microenvironmental influences.

Aging-associated immune cell dynamics also differed notably between males and females (**Fig. 4F**). For example, *Gzmk+* CD8 exhausted T cells exhibited greater expansion in females (**Fig. 4G, left**), notably in the lung, pancreas, and mWAT, while type 17 innate T cells showed a stronger increase in males, especially in the stomach, gWAT, and mWAT (**Fig. 4G, right)**.Overall, females exhibited broader immune expansion (53 female-biased vs. 31 male-biased subtypes), including *Tbx21+* aging-associated B cells (B cells-6), *Gpr55+ Ifi211⁺* NK cells (NK-3), *Sox5+ Ifnlr1+ Zfp362+* macrophages (Mac-11), and *Dtx1+ Srebp1a+* type II DCs (**Fig. 4H**), potentially contributing to women’s higher autoimmune prevalence (*60*). Meanwhile, we observed an enrichment of male-biased subtypes within gastrointestinal tissues (15/31 in male vs. only 2/53 in females) (**Fig. S13D**), such as the expansion of *Ccr9*+ ILC3s (NK/ILC-5) and *Cyp26b1*+ ILC3s (NK/ILC-6) in the intestine (**Fig. S13E**), reflecting more pronounced changes in the innate immunity during male intestinal aging (*61*).

We next extended analyses to non-immune cell lineages—vascular endothelial cells (VECs; n=426,160), fibroblasts (FB; n=377,181), and smooth muscle cells (SMCs; n=101,284) (**Fig. S15**). Unlike immune cells, these were strongly organ-specific (**Fig. S16**): 34/61 clusters were >90% derived from a single tissue, with tissue-specific TF programs identified. Examples include *Gata4*+ liver and *Foxf1*+ lung VECs (**Fig. S15C**), and tissue-specific TF programs in FBs (*Six1* bone marrow; *Foxf1* GI; *Tbx20* heart; *Hoxc9* kidney; *Tbx5* lung) (**Fig. S15F**). We also resolved female gonadal SMC subtypes marked by *Wt1/Esr1* (**Fig. S15I**).

These broadly distributed non-immune cells exhibited highly tissue-specific shifts in aging (**Fig. S15J-L, Table S6**). For example, we observed an age-associated depletion of endothelial cells, including cells in the kidney (VEC-14, *Igf1*+ kidney medulla endothelial), lung (VEC-7, *Tbx2*+ lung aerocytes), and muscle (VEC-1, *Aqp7*+) (**Fig. S17A**). By contrast, organ-specific endothelial cells in the liver (VEC-9, *Clec4g*+ sinusoidal endothelial) and pancreas (VEC-4, *Lamb1*+ VECs) expanded significantly during aging (**Fig. S17B**). The expansion of liver sinusoidal endothelial cells aligns with previous findings from *Tabula Muris Senis*(*48*). Fibroblasts (FB) likewise displayed tissue-specific aging trajectories, showing expansion in aged bone marrow (*Btla*+ FB-13), colon (*Edil3*+ FB-21), cecum (*Prkcq*+ FB-19), stomach (*Csf2rb*+ FB-3), and mWAT (*Nkx6-1*+ FB-17) (**Fig. S17C**). In contrast, two fibroblast subtypes (ovary *Tgfa*+ FB-9 and stomach *Sox6*+ FB-23) significantly decreased in aging (**Fig. S17D**). For smooth muscle cell (SMC) subtypes, there is a strong depletion of several subtypes across organs, such as colon (*Hoxd3*+ SMC-15), lung (*Tbx5*+ SMC-3), and ovary/uterus (*Gata2*+ SMC-2) (**Fig. S17F**). In the cecum, however, there was an increased population of SMCs marked by inflammatory gene markers such as *Il7* and *Serpina1c* (SMC-5, **Fig. S17G**).

Shared cross-organ patterns were identified, such as the depletion of a *Slc7a14*+ *Lhx6*+ *Meox2*+ vein endothelial cell subtype (VEC-5) (*56*, *57*) across several aged tissues (*e.g.,* muscle, gWAT, and ovary/uterus) (**Fig. S15M**), indicating widespread endothelial vulnerability. Conversely, an inflammatory FB subtype (*Ifi207/Ifi205/Ear1*+) expanded across mWAT, stomach, cecum, and esophagus in a male-specific manner (**Fig. S15N**). Re-analysis of published snRNA-seq atlas validated key trends, including the depletion of *Lhx6*+ endothelial cells in gWAT (**Fig. S15O)** and the expansion of *Csf2rb*+ fibroblasts in the stomach (**Fig. S15P**).

### Age- and sex-dependent epigenetic signatures of aging

We next investigated the organism-wide epigenetic aging program by examining 349 organ-specific main cell types using sex-stratified differential accessibility (DA) analyses, then linked DA sites to putative targets and inferred upstream regulators, including intrinsic transcription factors and extrinsic cytokine programs (**Methods, Fig. 5A**). This analysis revealed 1,318,007 (female) and 959,361 (male) aging-associated, cell type–specific DA peaks (**Fig. 5B, S18A-B**). Of 540,386 DA peaks shared between sexes in the same cell type, 96.9% changed concordantly (Spearman r=0.86 for logFC; p<2.2×10⁻¹⁶) (**Fig. 5C–D**).

**Figure 5.**
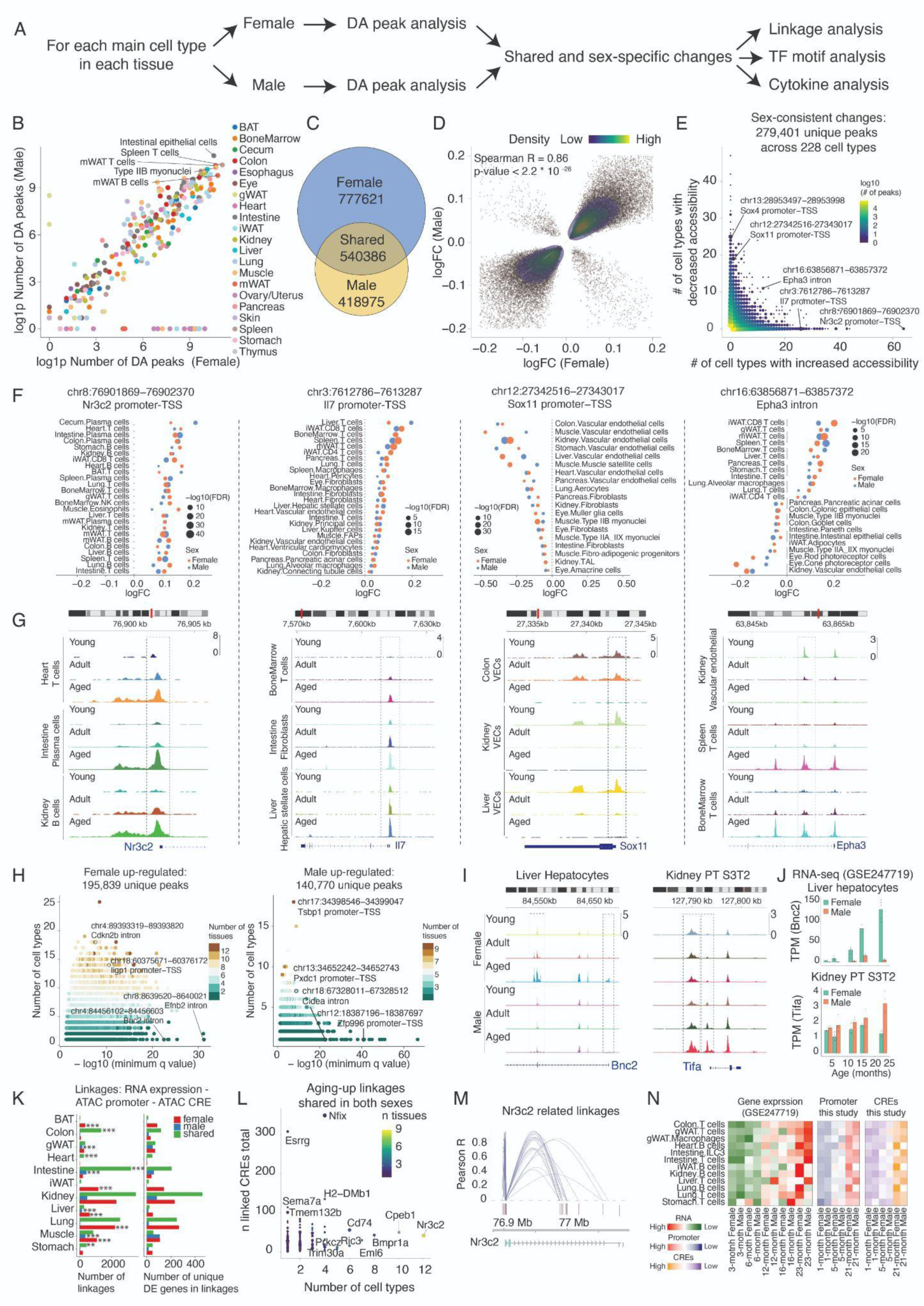
Aging-associated changes of chromatin accessibility. **(A)** Scheme for computational analyses to identify differentially accessible (DA) peaks along aging, their putative gene targets, and upstream regulators. **(B)** Scatterplot comparing the number of DA peaks between males and females for each main cell type. Cell types with a high number of DA peaks are labeled. **(C)** Venn plot of cell-type-specific DA peaks shared or unique to males and females. **(D)** Correlation of log transformed fold change (logFC, aging-associated accessibility changes per month, calculated via edgeR (*76*)) for DA peaks shared between sexes. **(E)** Scatterplot summarizing DA peaks with concordant changes with aging in both sexes. Axes show counts of cell types with increased (x-axis) or decreased (y-axis) accessibility; color denotes peak density. **(F)** Dot plots showing log transformed fold change (logFC) for DA peaks with consistent increases (promoters of *Nr3c2* and *Il7*), decreases (promoters of *Sox4*), or divergent trends (intronic peak of *Epha3*) across cell types. **(G)** Genomic tracks visualizing the accessibilities of DA peaks in (**F**) across three age groups. **(H)** Dot plots of sex-specific DA peaks for females (Left) and males (Right). Y-axis: number of cell types with significant increases; x-axis: -log10(q-value) for the most significant cell type. Color denotes the number of tissues with significant increases. **(I)** Genomic tracks of female-specific DA peak in liver hepatocytes (Left) and male-specific DA peak kidney proximal tubule cells S3T2 (Right). **(J)** Barplot showing validation of sex-specific chromatin accessibility changes using gene expression data (*7*) of the same genes in the same cell type as in (**I**). Dots represent gene expression of target genes across independent animals, quantified by transcripts per million (TPM). **(K)** Barplot showing the number of cell-type-specific, aging-associated gene-promoter-CRE linkages (Left) and the number of unique genes within those linkages (Right) for each tissue. Enrichment of aging-associated linkages in male-, female-, or shared-specific groups was evaluated per tissue by comparing observed counts with expectations derived from the global distribution across all tissues using a one-sided binomial test. Asterisks denote significance levels (**: q < 0.01; ***:q < 0.001). **(L)** Dot plot of genes with aging-upregulated linkages in both sexes. X-axis: cell-type count; y-axis: CRE count per gene. Color denotes the number of tissues containing cell types with significant increases. **(M)** Genome browser plot showing links between putative cis-regulatory elements and promoters for *Nr3c2*. **(N)** Heatmaps showing coordinated increases in gene expression, promoter accessibility, and linked distal CRE accessibility of *Nr3c2* across multiple immune cells with aging.

Most DA peaks were cell type–restricted (91.4% in ≤3 cell types), yet 803 peaks showed significant and consistent alterations across ≥10 organ-specific cell types (**Fig. 5E**), indicating coordinated aging programs. Notable examples include increased promoter accessibility of the mineralocorticoid receptor *Nr3c2* (63 cell types) and the proinflammatory cytokine *Il7* (26 cell types), predominantly within immune cells (**Fig. 5F–G**). Increased *Nr3c2* activity also marked the aging-associated B cell states (**Fig. 4B**). Conversely, we observed decreased promoter accessibility of *Sox4* (25 cell types) and *Sox11* (19 cell types), with Sox11 reductions prominent in endothelium (**Fig. 5E-G**). Some peaks changed in opposite directions by lineage (131 peaks with ≥5 cell types per direction). For example, an intronic *Epha3* peak increased in T cells/macrophages across 12 tissues but decreased in non-immune cells (**Fig. 5F–G, Fig. S18C**), underscoring context-dependent regulation.

We further examined the genomic distributions of aging-associated peaks. Both up-regulated and down-regulated peaks were enriched in intergenic regions relative to non-changing peaks (**Fig. S19A**). Notably, age-increased peaks showed strong enrichment within LINE and LTR elements, many of which harbor retrotransposons. Further analysis highlighted significant accessibility gains across fifty retrotransposon subfamilies, including MLT1F1, MLT1H, MMVL30-int, and RMER19B (**Fig. S19B-C**), consistent with reports of derepression of retrotransposons such as L1 elements in aging tissues (*8*). On the other hand, aging-decreased peaks had significantly higher conservation scores than stable peaks (**Fig. S20A**). This pattern was consistent across many individual tissues (**Fig. S20B**), indicating attenuation of conserved developmental regulatory programs with age.

Epigenetic remodeling in aging is highly sex-dependent. For example, we identified 195,893 female-specific and 140,770 male-specific sites with significantly increased accessibility in aging. Top female-specific DA peaks include intronic peaks at the senescence-associated gene *Cdkn2b* (in 18 cell types) and promoters of the interferon-inducible gene *Ligp1* (in 14 cell types) and *Ifi211* (in 16 cell types) (**Fig. 5H**). Cell–type–specific, sex-biased changes were evident: aged hepatocytes in females showed up-regulation of peaks overlapping with *Bnc2* (**Fig. 5I, left)**, a gene involved in extracellular matrix remodeling(*62*). In contrast, aged hepatocytes in males showed up-regulation of peaks overlapping with *Cidea* (**Fig. S18D, left**), a gene involved in lipid particle organization. In kidney PT S3T2 cells, *Mill2*-linked peaks were higher in aged females (**Fig. S18D**, right), while *Tifa* intronic peaks increased in aged males (programmed cell death) (**Fig. 5I**, right)—consistent with the male-specific decline of this state (**Fig. 2H**). These sex- and cell type–specific peaks matched promoter accessibility and were further confirmed by expression changes (**Fig. 5I–J**, **S18E**).

To connect CREs to their targets, we linked non-promoter DA peaks to nearby promoters via co-accessibility and then integrated a published snRNA-seq aging atlas (*7*) to identify genes exhibiting consistent promoter and expression changes with age. This yielded 21,360 (male) and 14,564 (female) cell type–specific gene–promoter–CRE linkages, with 12,862 shared across genders (**Fig. 5K-L**). Notably, *Nr3c2* was linked to 25 CREs, which showed increased accessibility during aging across multiple immune cell types and tissues, mirroring changes in promoter accessibility and gene expression from the same cell type (**Fig. 5M–N**).

### Intrinsic and extrinsic regulators of epigenetic reprogramming in mammalian aging

We then explored upstream regulators of these aging-associated chromatin changes by performing motif enrichment analyses of DA peaks in each cell type. Aging-upregulated peaks were enriched for inflammatory motifs that are confirmed across multiple cell types, including IFN-stimulated response elements (ISRE) in 80 cell types, IRF1 in 77 cell types, and AP-1 in 71 cell types (**Fig. 6A-B**). In contrast, downregulated peaks were enriched for TF families linked to stem cell maintenance (*e.g., SOX15* in 18 cell types) (**Fig. 6A**). Sex-specific patterns also emerged. For example, female-specific upregulated peaks were enriched for *POU* motifs in B cells, and male-specific upregulated peaks were enriched for inflammatory TF motifs (IRFs and AP-1) in intestinal epithelial cells (**Fig. 6C**).

**Figure 6.**
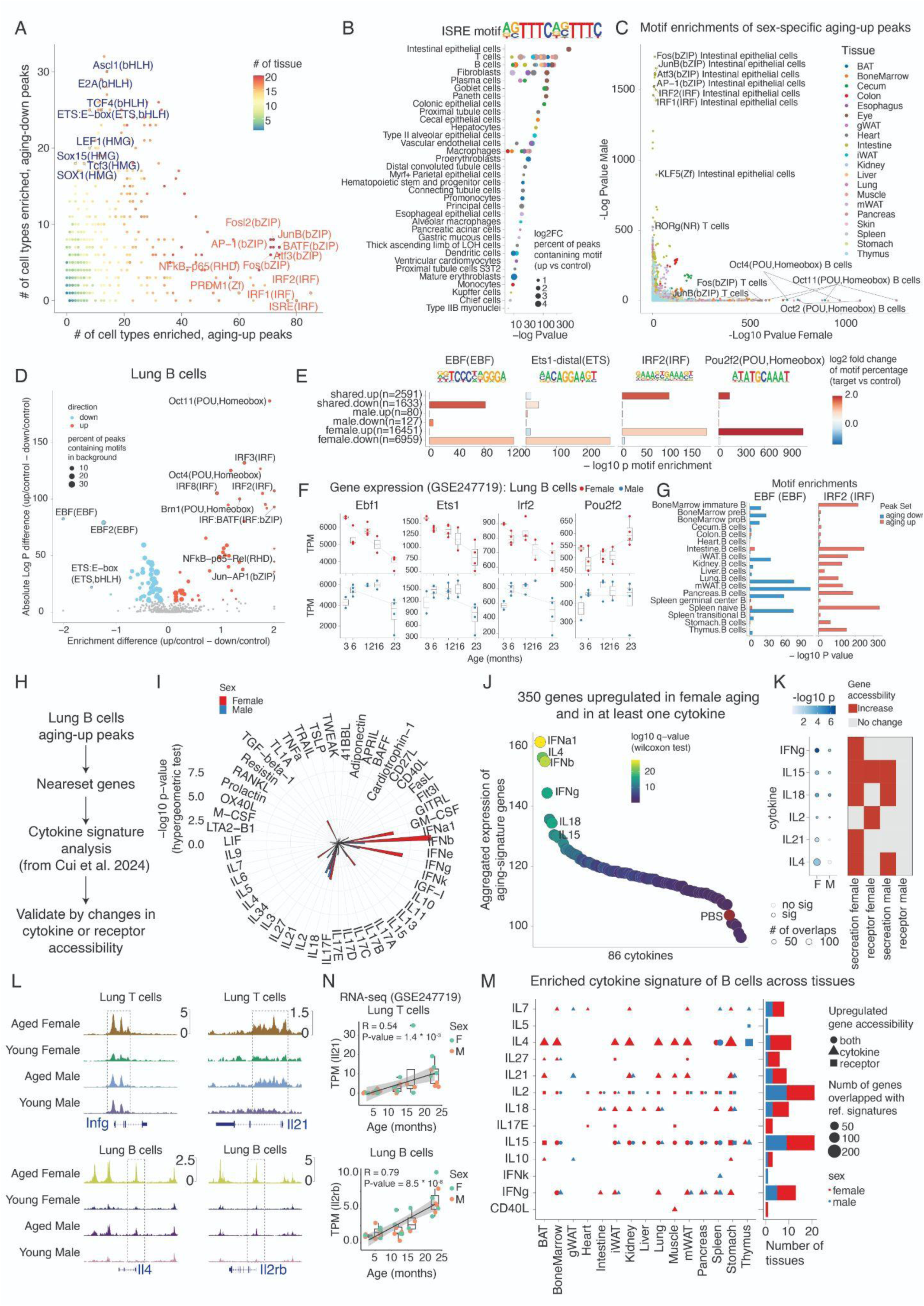
Identifying upstream regulators of aging-associated chromatin changes. **(A)** Sex-shared aging-associated up-regulated and down-regulated differentially accessible (DA) peaks for each main cell type were analyzed for transcription factor (TF) motif enrichment using HOMER (*77*). The scatter plot summarizes the number of cell types exhibiting significant motif enrichments (p.adj < 0.05) in either up-regulated (red) or down-regulated (blue) peak sets. TF motifs enriched across multiple cell types are labeled. **(B)** Dot plot showing the enrichment of interferon-stimulated response element (ISRE) motifs in aging-up DA peaks across cell types from diverse tissues. Dot size represents the fold change in ISRE-containing peaks in DA sets compared to a background peak set. Each dot represents one cell type from a specific tissue, colored by the tissue of origin. **(C)** Comparison of TF motif enrichment significance between female-specific and male-specific aging-up peak sets for each main cell type. Cell types with strong sex-specific motif enrichments are labeled. **(D)** Scatter plot showing enriched TF motifs in sex-shared, aging-up (red), and aging-down (blue) peaks from lung B cells. The x-axis denotes the difference in motif frequency (%) between DA and background peaks; the y-axis represents statistical significance. **(E)** Barplots showing motif enrichment for exemplary TFs: Ebf1 and Ets1 (down-regulated in both sexes), Irf2 (up-regulated in both sexes), and Pou2f2 (female-biased up-regulation) in lung B cells. **(F)** Scatterplot and barplot showing age-related expression changes of TFs highlighted in (**E**) in lung B cells, quantified using transcripts per million (TPM) for each animal. **(G)** Enrichment significance of EBF and IRF motifs for DA peaks from B cells across tissues, demonstrating conserved aging-associated regulatory patterns. **(H)** Scheme for computational analyses to identify cytokines potentially drive aging-associated chromatin remodeling in lung B cells. **(I)** Circle plot showing enrichments (-log10(p-value), hypergeometric test) of cytokine-specific signatures (derived from Immune Dictionary dataset (*67*)) in aging-associated genes identified in lung B cells from this study. **(J)** Dot plot displaying aggregated gene expressions of 350 genes shared between aging and cytokine treatments. Top conditions recapitulating aging-associated expression profiles were labeled. **(K)** Left: dot plot showing the enrichment score (-log10(p-value), hypergeometric test) of cytokine signatures in the male and female lung B cells as in **(I)**. Right: Heatmap indicating the aging-associated changes in the accessibility of cytokine secretion or receptor genes. **(L)** Genomic tracks showing aging-associated accessibility increases at *Ifng* and *Il21* in lung T cells and *Il4* and *Il2rb* in B cells. **(N)** Scatterplot showing increased gene expressions of *Il21* in T cells and *Il2rb* in B cells with age, together with a linear regression line. **(M)** Left, dot plot summarizing the enriched cytokine signatures using aging-up peak sets of B cells from each tissue. Only signatures that can be supported by increased accessibilities of secretion or receptor genes are shown. Right: barplot summarizing the number of tissues with activated signatures for each cytokine, splitting males and females.

These TF activity changes were further supported by the gene expression data. For example, in the aged B cells in the lung, we detected reduced motif accessibility for *Ebf1* and *Ets1* (**Fig. 6D-E**), both involved in B cell development (*23*) and quiescent state maintenance (*63*), aligning with their decreased expression in both sexes (**Fig. 6F**). Meanwhile, aged B cells exhibited decreased expression and increased motif accessibility for *Irf2* (**Fig. 6D-E**), reflecting its role as a transcriptional repressor of inflammatory signaling (*64*, *65*). Moreover, the aging-associated changes in EBF and IRF motifs were detected in B cells across tissues, highlighting a global chromatin transition in B cells during aging (**Fig. 6G**). Extending the analysis to other cell types, we examined target gene expression of aging-associated TFs using previously established gene–peak linkages (**Fig. S21**). Around one-quarter of TFs exhibited concordant changes between motif accessibility and expression of their linked targets within the same cell type, underscoring that epigenomic remodeling drives transcriptional reprogramming during aging.

We next explored the cooperative regulation by transcription factors during aging using a recent study that mapped TF–TF interaction motifs(*66*) (**Fig. S22A**). We identified 4,355 sex-shared aging-associated TF–TF pairs (**Fig. S22B and C**), most of which were concordant with at least one component TF. For example, the broad up-regulation of MAFF–FOSB in epithelial and immune cells was consistent with trends of each TF alone (**Fig. S22D**). Notably, some composite motifs exhibited stronger age-associated changes than each single factor, such as FOXA2–GABPA in gWAT monocytes and ELF1–FOXO1 in stomach T cells (**Fig. S22E**), suggesting epigenomic regulation during aging is shaped not only by individual TFs but also by composite TF interactions.

Given the prominent inflammatory signature, we next explored whether cytokines known to drive immune responses contribute to global epigenetic remodeling. Using the Immune Dictionary dataset (*67*), we compared aging-related DA peaks and associated genes with those induced by different cytokines (**Fig. 6H, Methods**). This revealed that aging signatures, especially in female lung B cells, resembled those induced by specific cytokines (*e.g.,* IFNs, IL4, and IL21) (**Fig. 6I**). Cross-referencing aging-upregulated gene expression changes with cytokine-responsive transcripts further validated these links (**Fig. 6J**). Concordant changes in chromatin accessibility of cytokine secretion and receptor genes further supported this link (**Fig. 6K-L**): For example, for secretion factors, we observed an increased accessibility of *Infg* and *Il21* in T cells, as well as *Il4* and *Il15* in B cells (**Fig. 6L**), aligning with activation of corresponding signatures, particularly in females (**Fig. 6I**). On the other hand, the upregulated response to IL2 and IL15 matched with increased accessibility of the receptor gene, *Il2rb*, in the aged B cells (**Fig. 6L**). A consistent trend was observed by analysis of recent snRNA-seq data (*7*), such as the increased expression of *Il21* in T cells, *Il2rb,* and *Il15* in B cells (**Fig. 6N** and **Fig. 6F**).

Extending the analysis across tissues, B cells showed widespread activation of cytokine programs with age (**Fig. 6M**). Notably, elevated IL21 signatures paralleled the global expansion of *Il21+* T cells (**T cells - 9, Fig. 4B** and **4D**) and were further confirmed at the human plasma protein level (*3*) (**Fig. 6G**). Prior work implicating IL21 in inducing Tbx21 and driving B-cell plasmacytic differentiation in SLE (*68*) suggests a similar role for IL21 in aging-related B-cell epigenomic remodeling. Applying the same framework to macrophages revealed activation of a distinct set—IL15, IL7, IL1α, IL18, and TNFα (**Fig. 6H**)—that may underlie age-related macrophage programs. Together, these data indicate global, sex-dependent epigenetic reprogramming linked to inflammatory signaling and cytokine-driven pathways.

## Discussion

Aging involves organism-wide cellular deterioration. While single-cell and spatial transcriptomics have mapped age-linked gene expression dynamics across cell types (*9*, *48*, *69*, *70*), systematic analyses of cell population shifts and the associated chromatin dynamics across the entire organism have been limited. Here, we profiled ∼7 million cells from 21 mouse tissues across three ages and both sexes to generate a single-cell chromatin accessibility atlas of aging. We reported the aging-associated dynamics of 536 tissue-level main cell types and 1,828 finer-grained subtypes, each defined by the accessibility across ∼1.3 million cis-regulatory elements. The dataset revealed widespread cell population shifts, lineage-specific epigenomic reprogramming, and striking sex-dependent patterns, laying a foundation for understanding the regulatory logic of mammalian aging.

We observed extensive cell population remodeling in aging: 146 of 536 organ-specific cell types (∼25%) changed significantly with age, including widespread increases of plasma cells and macrophages across various tissues, aligning with reported immunoglobulin accumulation (*70*) and monocyte/macrophage propagation (*71*) in aging. Non-immune vulnerabilities emerged across organs, such as the losses of kidney podocytes, muscle tenocytes and satellite cells, ovary granulosa cells, and lung aerocytes, likely contributing to losses of tissue homeostasis. High-resolution analyses of 1,828 subtypes showed nearly one-third with significant age dynamics, often diverging from their parent populations. Notable examples include sharp declines of naïve T-cell subtypes in lung and liver despite overall T-cell expansion, endothelial vulnerabilities specific to kidney medulla and vein, as well as a shared reactive epithelial state (*Creb5+ Bnc2+ Runx1+*) in kidney. Meanwhile, we identified shared cellular dynamics across organs, such as the synchronized expansion of many immune subtypes (*e.g., Gzmk+* CD8 T cells, regulatory T cells, *Il21+* CD4 T cells, *Il23r+* innate T cells, and *Tbx21+* B cells). This Ju also extends beyond mobile immunity, including depletion of *Slc7a14+/Lhx6+/Meox2+* vein endothelium and expansion of an *Ifi207+/Ear1+* aging-associated fibroblast program, indicating systemic inflammatory or hormonal cues underlying the synchronized cellular dynamics across organs.

At the molecular level, aging is accompanied by extensive chromatin reprogramming: 279,401 peaks changed with age, including a subset consistently altered across many cell types (*e.g.,* increased accessibility of *Nr3c2* and *Il7* promoters, *Sox4* or *Sox11* peaks that frequently lose accessibility). In addition, we observed increased accessibility of LINE and LTR elements as well as different retrotransposon subfamilies, including MLT1F1, MLT1H, MMVL30-int, and RMER19B. In contrast, the high conservation scores of aging-decreased peaks indicates down-regulation of conserved developmental regulatory programs. Motif analyses indicate a shift from stemness factors (*e.g., SOX, E2A*) to inflammatory regulators (*e.g., NF-κB, IRFs, AP-1*). Furthermore, integrating the data with a Immune Dictionary dataset (*67*) revealed specific cytokines (*e.g.,* IL15, IL21, IL4, and IFNγ) in driving the chromatin remodeling of aged B cells, implicating extrinsic drivers of cell-state transitions.

We found extensive sexual dimorphism in aging: ∼40% of aging-associated main cell types (55/146) and subtypes (193/499) showed sex differences. Overall, females display stronger immune activation at both compositional and regulatory levels, consistent with higher autoimmune prevalence in women (*72*, *73*). This included female-specific upregulation of TFs like *Pou2f2* in B cells, matching female-biased expansions of aging-associated B cell subtypes. Some subtypes shift in opposite directions by sex, such as *Slc22a7*+ proximal tubule cells (PT S3T2) increasing in aged females but decreasing in males. At the molecular level, tens of thousands of cell-type–specific peaks change in only one sex, underscoring sex as a major axis of aging heterogeneity.

This study has several limitations. First, our cell analysis is based on mice and thus its direct relevance to human biology requires further comparative analyses. In addition, this study focuses on chromatin accessibility from dissociated nuclei, yet this inevitably misses orthogonal molecular (*e.g.,* RNA, protein) and spatial information that could further elucidate the functional consequences of regulatory changes. Additional modalities and time points would further refine aging trajectories and mechanisms.

Despite these limitations, our dataset provides a valuable framework for exploring the molecular basis of human aging and disease. Many cis-regulatory elements identified here overlap with orthologous regions in the human genome, enabling cross-species mapping of regulatory programs. Integrating aging-associated regions with human GWAS loci will help prioritize cell types and regulatory elements most likely to mediate disease risk. Moreover, the systematic characterization of sex-dimorphic cell states in our atlas offers insights into sex-biased disease prevalence, including autoimmune disorders and kidney disease. Together, these features position our dataset as a resource for understanding the molecular logic of mammalian aging and for guiding therapeutic strategies aimed at preserving or restoring youthful tissue states.

## Acknowledgments

We thank members of the Cao lab for helpful discussions and feedback. We also thank members of the Rockefeller University High-Performance Computing Core, Comparative Bioscience Center, Genomics Resource Center, Bioinformatics Resource Center and Flow Cytometry Resource Center for their invaluable support.

## Author contributions

J.C. and W.Z. conceptualized and supervised the project; Z.L. optimized nuclei extraction methods and EasySci-ATAC protocols; Z.L. performed mouse dissection, nuclei extraction, and single-cell ATAC-seq experiment; Z.L. performed all computational analyses with insights from Z.X., Z.Z., and A.A.; J.C., W.Z., and Z.L. wrote the manuscript with input from all co-authors.

## Data availability

Raw FASTQ files, processed count matrices, cell metadata, and peak metadata can be downloaded from NCBI GEO under accession number GSE288730. An interactive website facilitating the visualization of our data across cell type, age, and sex is available at https://epiage.net.

## Code availability

Custom code and scripts used for processing sequencing reads of the EasySci-ATAC-seq library were deposited in Zenodo (*74*) (https://doi.org/10.5281/zenodo.8395492).

## Supplementary Materials

## Materials and Methods

### Animals and organ collection

The C57BL/6 wild-type mice at one month (n=10, 5 males and 5 females), five months (n=10, 5 males and 5 females), and twenty-one months (n=12, 6 males and 6 females) were obtained from The Jackson Laboratory. Detailed information on animal individuals in this study is provided in Table S1A. Most libraries pooled all animals to minimize batch effects between samples. Among the 21 tissue types, 10 were completed in a single experiment (and therefore correspond to a single library), while the remaining tissues were completed across 2–4 experiments. The experiment IDs have been included in the metadata table of the submitted processed dataset. Mice were housed socially and maintained on a regular 12h/12h day/night cycle. In order to analyze newborn cells from the same set of samples in parallel, all mice were labeled with EdU (25 mg/kg, i.p. injection) at 24-hour intervals for five days. Tissues were harvested one day after the final injection. To control for circadian effects, sample harvest was performed around the same period (4-7 PM) across all individuals. Mice were euthanized utilizing inhalation of carbon dioxide (CO2). After euthanization, major organs were quickly transferred into ice-cold PBS, and the following tissues were collected: brown adipose tissue, bone marrow, cecum, colon, esophagus, eye, heart, small intestine (spanning duodenum, jejunum, and ileum), kidney, liver, lung, muscle (quadriceps and gastrocnemius), ovary and uterus, pancreas, back skin, spleen, stomach, thymus, gonadal white adipose tissue, inguinal white adipose tissue, mesenteric adipose tissue. Dissected mouse tissues were snap-frozen in liquid nitrogen and stored at −80C. All animal procedures were in accordance with institutional regulations under the IACUC protocol 21049.

### Nuclei extraction from multiple mammalian organs

Nuclei extraction was performed using the method from (*78*) and (*7*) with slight modifications. Before extraction, frozen tissues were placed inside aluminum foil, and smashed into powders on dry ice with a hammer. 10X PBS-hypotonic stock solution was prepared as follows: 6.83 g of Na2HPO4-2H2O (Sigma, 71643-250G), 3.5 g of NaH2PO4-H2O (CATALOG), 1.2 g of KH2PO4 (Sigma, P285-500), 1 g of KCl (Sigma, P9541-1KG) and 3 g of NaCl (Sigma, P9888-500G) in nuclease-free water to a final volume of 500 mL. On the day of nuclei extraction, 1X nuclei lysis buffer was prepared freshly as follows: final concentration of 1X PBS-hypotonic stock solution, 3 mM MgCl2 (VWR, 97062-848), 0.025% IGEPAL CA-630 (VWR, IC0219859650), 0.1% Tween-20 (Sigma, P9416-100ML), 1X cOmplete, EDTA-free Protease Inhibitor Cocktail (Sigma, 11873580001). In addition, a final concentration of 0.33M sucrose (Sigma, S0389) was included in the lysis buffer for these tissues: esophagus, stomach, intestinal, cecum, colon, spleen, thymus, bone marrow and skin. The powdered tissues were then transferred into 10-20 mL nuclei lysis buffer, followed by brief vortexing and incubation for 10-15 minutes at 4C on a rotator. Extracted nuclei were then filtered through 40 um cell strainers (VWR, 470236-276), stained with 4’,6-diamidino-2-phenylindole staining (DAPI, Invitrogen D1306), and FACS sorted for singlets.

### EasySci library construction and sequencing

The EasySci-ATAC library was prepared following the prior study (*9*), except for a few modifications. The sorted nuclei were pelleted down at 500g for 5 minutes, resuspend in nuclei buffer [10 mM Tris-HCl pH 7.5 (VWR, 97062-936), 10 mM NaCl (VWR, 97062-858), 3 mM MgCl2 (VWR, 97062-848), 0.025% IGEPAL CA-630 (VWR, IC0219859650), 0.1% Tween-20 (Sigma, P9416-100ML), 1X cOmplete, EDTA-free Protease Inhibitor Cocktail (Sigma, 11873580001)] to a final concentrations of 1000∼2000 nuclei/µL. Nuclei were mixed in a 1:1 ratio with 2X TD buffer [20 mM Tris-HCl pH 7.5, 20 mM MgCl2, 20% Dimethylformamide (Fisher, AC327175000)] and dispensed 10 µL into each well of four 96-well plates. 1 µL barcoded Tn5 was loaded into each well. Tagmentation reaction was performed at 37°C for 30 minutes with gentle shaking at 300 rpm and stopped by adding 11 µL of 2X Stop buffer [40 mM EDTA (VWR, 37062-656), 1 mM Spermidine (Sigma, S0266-1G)] to each well. Samples were pooled and washed twice and resuspended in nuclei buffer. 5 µL of resuspended nuclei were distributed into each well of 96-well plates. 2 µL indexed EasySci P5 ligation adapters and 3 µL ligation mix [1 µL nuclease-free water, 1 µL 10X T4 DNA ligase buffer, 1 µL T4 DNA ligase (NEB, M0202L)] were added to each well. Ligation was performed at room temperature for 30 minutes with medium-speed rocking (350g) and stopped by adding 2 µL of 18 mM EDTA to each well. After that, nuclei were pooled, washed, resuspended using nuclei buffer, and subjected to another round of FAC sort based on DAPI staining to remove doublets. Then, sorted nuclei were distributed into PCR plates as 5 uL per well. Proteinase K treatment was performed by mixing each well with 0.25 μL 18.9 mg/mL proteinase K (Sigma, 3115828001), 0.25 µL 1% SDS, and 0.5 µL EB buffer, and plates were incubated at 65°C for 16 hours. Then, 2 μL 10% Tween-20 was added to each well to quench the SDS. Following on, 1 μL of 10 μM universal P5 primer (5′-AATGATACGGCGACCACCGAGATCTACAC-3′, IDT), 1 μL of 10 μM indexed P7 primer (5’-CAAGCAGAAGACGGCATACGAGAT[i7]GTGACTGGAGTTCAGACGTGTGCTCTTCCGATCT-3’, IDT) and 10 μL NEBNext High-Fidelity 2X PCR Master Mix (NEB M0541L) were added into each well. Amplification was carried out using the following program: 72°C for 5 minutes, 98°C for 30 seconds, 13-14 cycles of 98°C for 10 seconds, 66°C for 30 seconds, 72°C for 30 seconds and a final 72°C for 5 minutes. Final PCR products were pooled and purified by column purification using a Zymo DNA Clean & Concentrator kit (Zymoresearch, D4014) followed by gel extraction using Zymoclean Gel DNA Recovery Kit (Zymoresearch, D4007) to remove adapter dimers. Library concentrations were determined by Qubit and the libraries were visualized by electrophoresis on a 2% E-Gel™ EX Agarose Gels (Invitrogen G402022).

### Processing of sequencing reads

Base calls were converted to fastq format and demultiplexed using Illumina’s bcl2fastq/v2.19.0.316 tolerating one mismatched base in barcodes (edit distance (ED) <= 1). Then, indexed Tn5 barcodes and ligation barcodes were extracted, and corrected to their nearest barcode (edit distance (ED) <= 1). Reads with uncorrected barcodes (ED >= 2) were removed. Tn5 adaptors were removed from 5’-end and clipped from 3’-end using trim_galore/0.4.1 (https://github.com/FelixKrueger/TrimGalore). Trimmed reads were mapped to the mouse genome (mm10) using STAR/v2.5.2b with default settings. Aligned reads were filtered using samtools/v1.4.1 to retain reads mapped in proper pairs with quality score MAPQ > 30 and to keep only the primary alignment. Duplicates were removed by picard MarkDuplicates/v2.25.2 per PCR sample. Deduplicated bam files were converted to bedpe format using bedtools/v2.30.0, which were further converted to offset-adjusted (+4 bp for plus strand and −5 bp for minus) fragment files (.bed). Deduplicated reads were further split into constituent cellular indices using the Tn5 and ligation barcodes, and sparse matrices counting reads overlapping with promoter regions (±1 kb around transcription start site) were generated for quality filtering. In the meantime, fragment files were used to generate h5ad files for all downstream analyses with the snap.pp.import_data() function of SnapATAC2/v2.5.1. Cell-by-bin matrices were also generated with SnapATAC2 function snap.pp.add_tile_matrix() containing insertion counts across genome-wide 5000-bp bins.

### Cell filtering, dimensionality reduction, clustering, and annotations

The following analyses were performed on the dataset of each tissue separately. After initial processing, cells with less than 1000 unique reads or less than 15% of reads in promoter regions were discarded. The promoter ratio cutoff was adjusted to 0.1 for the eye dataset due to the observation of a lower promoter ratio in corneal epithelial cells. Then, we used an iterative clustering strategy to detect potential doublet cells from each organ, similar to our previous study (*9*, *15*). Briefly, doublet scores were calculated for each cell using SnapATAC2 function snap.pp.scrublet(). Meanwhile, all cells of each organ were subjected to clustering and sub-clustering analysis with spectral embedding and graph-based clustering implemented in SnapATAC2. Cells labeled as doublets (defined by a doublet probability cutoff of 0.5) or from doublet-derived sub-clusters (defined by a doublet ratio cutoff of 0.2) were manually examined and filtered out. We then generated gene activity matrices for each organ by counting the Tn5 insertions in the TSS and gene body regions for each gene using SnapATAC2 function snap.pp.make_gene_matrix(). Gene activity matrices were then used for cell type annotations.

To identify clusters of cells corresponding to different cell types, we subjected cells after data cleaning to dimension reduction and Leiden clustering using SnapATAC2 functions snap.tl.spectral(), snap.pp.knn() and snap.tl.leiden() with the default setting. UMAP coordinates were calculated based on the spectral embedding matrices using the function UMAP() with min_dist=0.01 from the Python package umap/v0.5.2 (*79*). For cell annotations, we first obtained a draft of annotations by integrating our chromatin data (subsampled to 2,000 cells per Leiden cluster) with the published sn-RNA-seq datasets (*7*) through Seurat/4.3.0.1 (*7*, *80*) label transfer, and we manually reviewed and refined the annotations for each Leiden cluster based on accessibilities of known markers listed in Table S2.

Reproducibility of cell-type annotations was assessed using leave-one-out cross-validation with a support vector machine classifier, as described previously (*15*). For each tissue, cells from all but one animal were used to train SVM classifiers on spectral embedding matrices, and the trained model was applied to predict cell type labels in the hold-out animal. This procedure was repeated across all biological replicates, and reproducibility was quantified as the overall prediction accuracy on all out-of-fold data.

### Peak calling

After cell annotation, reads from each main cell type of each tissue were concatenated. Then, Tn5-corrected single-base insertions were extracted and subjected to peak calling using macs3/v3.0.0b3 (*81*), with the following parameters: --nomodel --extsize 200 --shift −100 -q 0.1. Peak summits were extended by 250bp on either side and then merged iteratively, similar to (*18*). Specifically, the peak with the smallest p-value was kept and any peak that directly overlapped with it was removed. Then, this process was repeated to the second most significant peak and so on until all peaks were either kept or removed. Interactive merging was performed first across main cell types within each tissue, and then across all tissues to generate the universal peak set across the entire organism. The peak count matrices were generated using the SnapATAC2 function snap.pp.make_peak_matrix(). Peak annotations were performed using the HOMER function annotatePeaks.pl.

### Identification of cell-type-specific peaks

We used a Shannon entropy-based method to identify cell type-specific peaks, similar to previous studies (*18*, *82*). Starting from 144 unique cell types collapsed from all tissues, we first aggregated the peak count matrix for each cell type and normalized the aggregated matrix to counts per million (CPM). We then converted the CPM values into probabilities by dividing each cell type’s CPM by the total CPM across all cell types for a given peak: p_i = q_i / Σ q_i, where “q_i” is the CPM value of the i-th cell type of a given peak. Next, we calculated Shannon entropy for each peak, which measures how uniformly accessibility is distributed across cell types: H = - Σ (p_i * log(p_i)) where “p_i” is the probability of the i-th cell type for a given peak. To determine statistical significance, we generated a background distribution of entropy values using a set of peaks with low variability. Specifically, for each peak, we calculated the fold change between the most accessible cell type (quantified by CPM) and the average accessibility across all cell types. We then grouped peaks based on their mean accessibility into thirty bins. Within each bin, peaks with fold-change values below the 25th percentile were considered low-variable peaks, while those in the top 25th percentile were considered high-variable peaks. The entropy threshold was set at a p-value of 0.05, using the low-variability peaks as the background. Cell type-specific peaks were identified from the highly variable peak set if their entropy values fell below this threshold. Cell-type-specific peaks, along with their maximum and average accessibility values quantified by CPM, can be found in Table S3.

### LDSC analysis

To estimate enrichments of heritability for human traits in cell-type-specific peaks, we applied LDSC (*83*), which takes summary statistics from a given GWAS as input and quantifies the enrichment of heritability in an annotated set of SNPs conditioned on a baseline model that accounts for the non-random distribution of heritability across the genome. The LDSC computational pipeline was modified from (*12*) and the LDSC tutorial (https://github.com/bulik/ldsc/wiki/Cell-type-specific-analyses). Specifically, we first used the UCSC utility liftOver to lift all GWAS SNPs from the human to the mouse genome. We then took the top 2000 differentially accessible peaks per main cell type identified from the entropy-based method, and annotated each SNP according to whether or not it overlapped these cell-type-specific peaks. We then followed the recommended workflow for running LDSC using HapMap SNPs, precomputed files corresponding to 1000 genomes phase 3, excluding the MHC region to generate an LDSC model for each chromosome and peak set. GWAS summary statistics were obtained from the Broad LD Hub (https://data.broadinstitute.org/alkesgroup/sumstats_formatted/) and from GWAS catalogs (*84*) (https://www.ebi.ac.uk/gwas/downloads/summary-statistics). Coefficient P values calculated from LDSC were corrected for multiple hypotheses for each trait using the Benjamini-Hochberg method.

### Gene-peak linkage analysis

RNA–ATAC linkage analysis was performed based on gene–peak correlations across matched cell types. Specifically, within each age group, we collapsed single-cell peak-count and gene-count matrices by tissue and cell type to generate matched pseudo-bulk profiles. For each gene, we then computed the Pearson correlation between its expression and the accessibility of all nearby peaks (±100 kb of the TSS) across cell types. To assess significance, we generated a background distribution by shuffling cell-type labels in the aggregated scRNA-seq profiles and determined a correlation threshold controlling the false discovery rate at 0.05.

### Identification of sex dimorphism for the main cell types

To examine cell types with sex dimorphism in chromatin accessibility profiles, we trained k-nearest neighbor (KNN) classifiers to distinguish male and female cells of the same age for each main cell type using python package sklearn/v1.0.2. Spectral embeddings generated from SnapATAC2 were used as input features to predict the sex of individual cells. Datasets were filtered to include only cell types with at least 200 total cells and a minimum of 50 cells per sex. We performed five independent train-test splits, using 80% of the data for training and 20% for testing, with stratified sampling to maintain sex balance. To ensure computational efficiency, we subsampled both training and test sets to a maximum of 2000 cells per cell type. Classification performance was evaluated using area under the curve (AUC) metrics, with cell types exhibiting AUC > 0.9 considered to display strong sex-associated chromatin differences.

### Sub-clustering analysis

Sub-clustering analysis was performed using SnapATAC2, same as main clustering. Briefly, we subjected cells of the same type within the same tissue to feature selection (n_features=50000), dimension reduction and leiden clustering (resolution=1.5). UMAP coordinates were calculated based on the spectral embedding matrices (min_dist=0.01). We then examined the resulting Leiden clusters and manually merged those that overlapped in UMAP space without clear differences in gene accessibility. For certain cell types with pronounced sex dimorphism, including liver hepatocytes, kidney proximal tubule cells, type IIB myonuclei in muscle, adipocytes, and adipose stem and progenitor cells from gonadal white adipose tissue, sub-clustering was performed separately for males and females to capture cell-state differences rather than sex-based differences.

### Differential Abundance Analysis

For each mouse individual, we quantified the proportion of each main cell type or subtype within each tissue, and applied a linear regression model (proportion ∼ age + age:sex) to assess age-associated population dynamics while accounting for sex effects. The analysis was conducted using the R function lm(). Quadriceps and gastrocnemius muscles were analyzed separately, as they were from distinct anatomical locations. In the model, age was treated as a continuous variable (in months), while sex was considered a binary categorical variable. To identify significantly changing subtypes, we filtered results based on the following criteria: R² > 0.4 and a q-value (for either the age term or the age:sex interaction term) < 0.05. Multiple hypothesis correction was performed separately for each tissue using the Benjamini-Hochberg method. Additionally, Pearson correlation coefficients were calculated between the proportion of each main cell type or subtype and age. Based on these correlations, cell types and subtypes were classified into three groups: aging-up (q-value of the age term < 0.05, q-value of the age:sex interaction > 0.05, Pearson r > 0), aging-down (q-value of the age term < 0.05, q-value of the age:sex interaction > 0.05, Pearson r < 0), interaction-significant (q-value of the age:sex interaction < 0.05).

### Integration analysis between RNA-seq and ATAC-seq

Integration analysis of main cell types was performed using a standard Seurat anchor-based workflow. For each tissue, 10,000 cells per main cell type per modality were subsampled. Anchors were selected using FindTransferAnchors() functions, and two datasets were then integrated together and co-embedded to the same low-dimensional space.

For subtype-level comparison, scBridge (*58*) was used to transfer subtype annotations from scATAC-seq to scRNA-seq data of the same main cell types. Subtype validity was then assessed using two orthogonal criteria: molecular similarity and population dynamics consistency.For molecular similarity, the top 50 ATAC-derived marker genes per subtype were identified using scanpy.tl.rank_genes_groups and ranked by log fold-change. Cross-modality correlations were then computed between matched and unmatched subtypes within the same main cell type. For population dynamics consistency, differential abundance analysis was performed on RNA-seq cells using the same procedure as for ATAC-seq, and the directionality of subtype changes across modalities was compared.

### Differential Peak Analysis

In order to reduce the search space for differential peak analysis, we first selected highly accessible and highly variable peaks for each main cell type. First, the accessibility levels of all 1.3M peaks for each condition were quantified by counts per million (CPM) after aggregating data of the same age. Peaks were then ranked based on their CPM value in the most accessible condition, and only those above the 75th percentile were retained for each main cell type. These highly accessible peaks were further grouped into thirty bins based on their maximum accessibility. Within each bin, peaks with fold-change values (max condition vs. mean of three age groups) in the top 25th percentile were classified as highly variable peaks. This preselection was conducted separately for males and females, and the resulting peaks were collapsed across sexes.

Pre-filtered peaks for each main cell type were then analyzed for differential accessibility using edgeR/v3.36.0 (*18*, *76*). The analysis was performed at the pseudo-bulk level by aggregating single-cell peak counts from the same mouse sample, treating each individual mouse as a replicate. Age was treated as a continuous variable (in months). Only cell types with more than 200 cells in all three age groups were included in the analysis. Additionally, male and female samples were tested separately, with log fold-change (logFC), p-values, and q-values computed for each sex independently using edgeR. Differentially accessible peaks were categorized as follows: sex-shared, aging up-regulated peaks (p-value-female < 0.05 and p-value-male < 0.05, q-value-male < 0.05 or q-value-male < 0.05, logFC-female > 0, logFC-male > 0); sex-shared, aging down-regulated peaks (p-value-female < 0.05 and p-value-male < 0.05, q-value-male < 0.05 or q-value-male < 0.05, logFC-female < 0, logFC-male < 0); female-specific DA peaks (p-value-male > 0.05, q-value-female < 0.05); male-specific DA peaks (p-value-female > 0.05, q-value-male < 0.05).

### Identifying aging-associated linkages of genes, promoters and cis-regulatory elements

This analysis aims to identify links between cis-regulatory elements and their putative target genes, and to validate accessibility changes with gene expression data. First, we focused on aging-associated differentially accessible (DA) peaks identified previously. Pearson correlation coefficients were computed between DA promoters and nearby non-promoter DA peaks (±500 kb) across samples, after pseudo-bulking each mouse individual. A background distribution was also generated by pairing DA promoters with randomly selected peaks, and a threshold of Pearson correlation was defined with an empirical FDR = 0.05. The correlation analyses were performed separately for sex-shared, female-specific and male-specific peaks. In the meantime, we collected snRNA-seq data of mouse aging from (*7*), re-annotated the cells for consistent cell type labels between RNA-seq and ATAC-seq, and followed with differential gene expression changes using edgeR/v3.36.0 (*76*). Similar to differential accessible analysis, the differential expression analysis was performed separately for males and females in terms of each main cell type in each tissue, and at the pseudo-bulk level by aggregating single-cell gene counts from the same mouse individual. Differentially expressed genes, including sex-shared (p-value < 0.1 in both males and females, with the same logFC direction), female-specific (p-value < 0.1 in females but > 0.1 in males) and male-specific changes (p-value < 0.1 in males but > 0.1 in females) were defined. Finally, aging-associated linkages (gene-promoter-CRE) were defined if a gene exhibited consistent changes (up-regulated or down-regulated) in the same cell type from the same tissue across three layers, i.e, expression changes based on RNA-seq data, accessibility changes of promoters and non-promoter peaks based on ATAC-seq data, significant correlations between accessibilities of promoters and linked non-promoter sites.

### Transcription factor motif analysis

We used Signac/v1.7.0 (*85*) and chromVAR/v1.16.0 (*19*) to quantify motif accessibility of transcription factors in Figure 3N, 5C, 5I, S3E, S3F and S7B. Position weight matrices of transcription factor binding sites were obtained from JASPAR2022 (*86*). A Signac object containing a peak count matrix of cells of interest was constructed using the Signac functions CreateChromatinAssay() and CreateSeuratObject(). Motif information was then added via AddMotifs(), and motif deviation scores were calculated for each single cell using RunChromVAR(). In Figure S3F, to compare motif activity and gene accessibility of transcription factors across cell types, motif deviation scores were rescaled to the range (0,10) using the R function rescale(), averaged per cell type, and scaled to z-score across cell types. In Figure 5C, motif footprinting analysis was performed using the Footprint() function in Signac.

For motif enrichment analysis of aging-associated differentially accessible (DA) peaks (Figure 7A, 7C, 7D, and 7E), we used the HOMER function findMotifsGenome.pl (*77*). A background peak set with minimal age-related changes was selected for each main cell type as a control for HOMER analysis. Specifically, accessible peaks for each cell type were grouped into thirty bins, based on accessibility in the highest condition. Within each bin, peaks with fold-change values (maximum condition vs. the mean of three age groups) in the bottom 10th percentile were classified as invariable peaks. The same background peak set was used to assess transcription factor motif enrichment across all aging-associated peak sets of a given cell type. Motif enrichments were quantified as the percentage of peaks containing a given motif in the target peak set relative to the background. Significance values (-log10 q-value) were calculated within HOMER.

Combinatorial effects of transcription factors during aging were assessed using a published resource of cooperative binding motifs for TF pairs (*66*). For each main cell type, sex-shared aging-associated differentially accessible peaks were extracted, and TF–TF interaction motifs were mapped to these peaks using the provided position weight matrices. A matrix of motif-by-cell counts was then obtained by multiplying the peak-by-cell matrix with the peak-by-motif matrix, and was aggregated for each biological replicates. Differential motif accessibility analyses were then performed using edgeR (*76*) across ages for each main cell type, and significant results were determined based on an FDR cutoff of 0.05.

### Analysis of TF target gene expression

Gene–peak linkages (Fig. R1-3) were used to examine expression changes in targets of age-influenced transcription factors. For each cell type, we identified aging-associated peaks containing binding sites of age-influenced TFs using HOMER and used the gene-peak linkages to define each TF’s target genes. Pseudo-bulk expression profiles per animal were generated from the PanSci scRNA-seq dataset. Aggregated normalized expression of TF target genes was then compared between 6- and 23-month-old mice using Wilcoxon rank-sum tests to determine significant changes on the expression of target genes (p < 0.05).

### Cytokine-related analysis

To evaluate the contribution of cytokine signaling to age-related molecular changes in B cells and macrophages, aging-upregulated DA peaks of each sex identified from chromatin accessibility data were first collapsed to their nearest genes using the HOMER function annotatePeaks.pl. The resulting gene sets (female-upregulated and male-upregulated) were compared to cytokine-induced gene sets of the same cell type from the Immune Dictionary Data (*67*). A hypergeometric test was performed using the R function phyper(). We only kept conditions containing more than ten DE genes after cytokine treatments identified from the Immune Dictionary for this comparison.

As an additional measure of aging signatures across cytokine treatments, we overlapped the aging-upregulated gene signatures from our study with the union of genes that exhibited increased expression in at least one cytokine treatment. This overlapping gene set represents shared features between aging and cytokine responses. To identify cytokines that mimic aging phenotypes, we compared the aggregated expression of these genes across cytokine treatments to the PBS control. Statistical evaluation was conducted using Wilcoxon rank-sum tests (wilcox.test() in R).

To minimize cross-activation effects, we excluded cytokines whose receptors were not expressed (TPM < 1) in the target cell type, based on gene expression profiles from the Immune Dictionary. To further validate cytokine signaling changes and refine potential targets, we further analyzed chromatin accessibility changes in genes encoding cytokines and their receptors. Differential analyses were performed using gene accessibility matrices at the pseudo-bulk level, with females and males analyzed separately.

Finally, cytokine signatures were defined based on the following criteria: significant overlap with aging-peak-associated genes (p-value < 0.05, hypergeometric test); significant aging-mimicking activation compared to PBS (p-value < 0.05, Wilcoxon rank-sum test); can be supported by increased chromatin accessibility of the underlying genes in at least one cell type within the same tissue, or from increased chromatin accessibility of receptor genes in the target cell type.

## Supplementary figures

**Figure S1.**
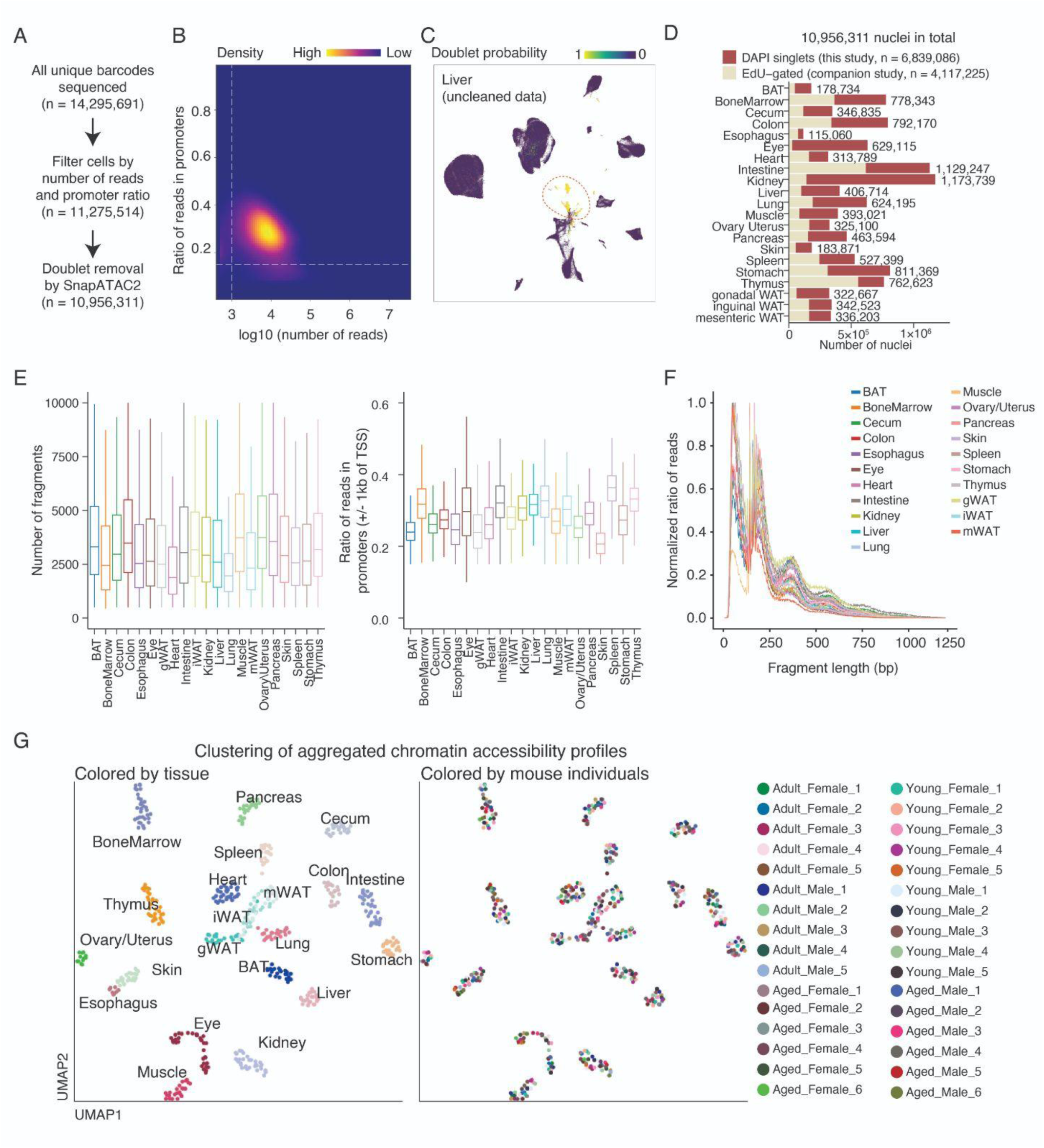
Overview of dataset quality. (A) Scheme of data cleaning procedures. All sequenced cells underwent initial filtering determined by reads number and promoter ratio, followed by doublet removal using a modified pipeline based on SnapATAC2 (*14*). (B) Density plot showing the distribution of the number of reads per nucleus versus the ratio of reads in promoters. Dash lines indicated cutoffs used for filtering (promoter ratio > 0.15 and > 1,000 unique reads). (C) An example UMAP plot showing all liver cells (before doublet removal), colored by doublet probability. Circle indicated doublet cells. (D) Barplot showing the number of cells profiled in each tissue, colored by the sources. DAPI singlets represent the global cell population investigated in this study. (E) Box plot showing the number of fragments (left) and the ratio of reads mapped to promoters (± 1kb of TSS, right) per nucleus across tissues. (F) Line plot showing the fragment length distributions of aggregated single cell ATAC-seq data across tissues. (G) UMAP visualization of the aggregated chromatin accessibility profiles from all samples across tissues, colored by organ (left) and mouse individuals (right).

**Figure S2.**
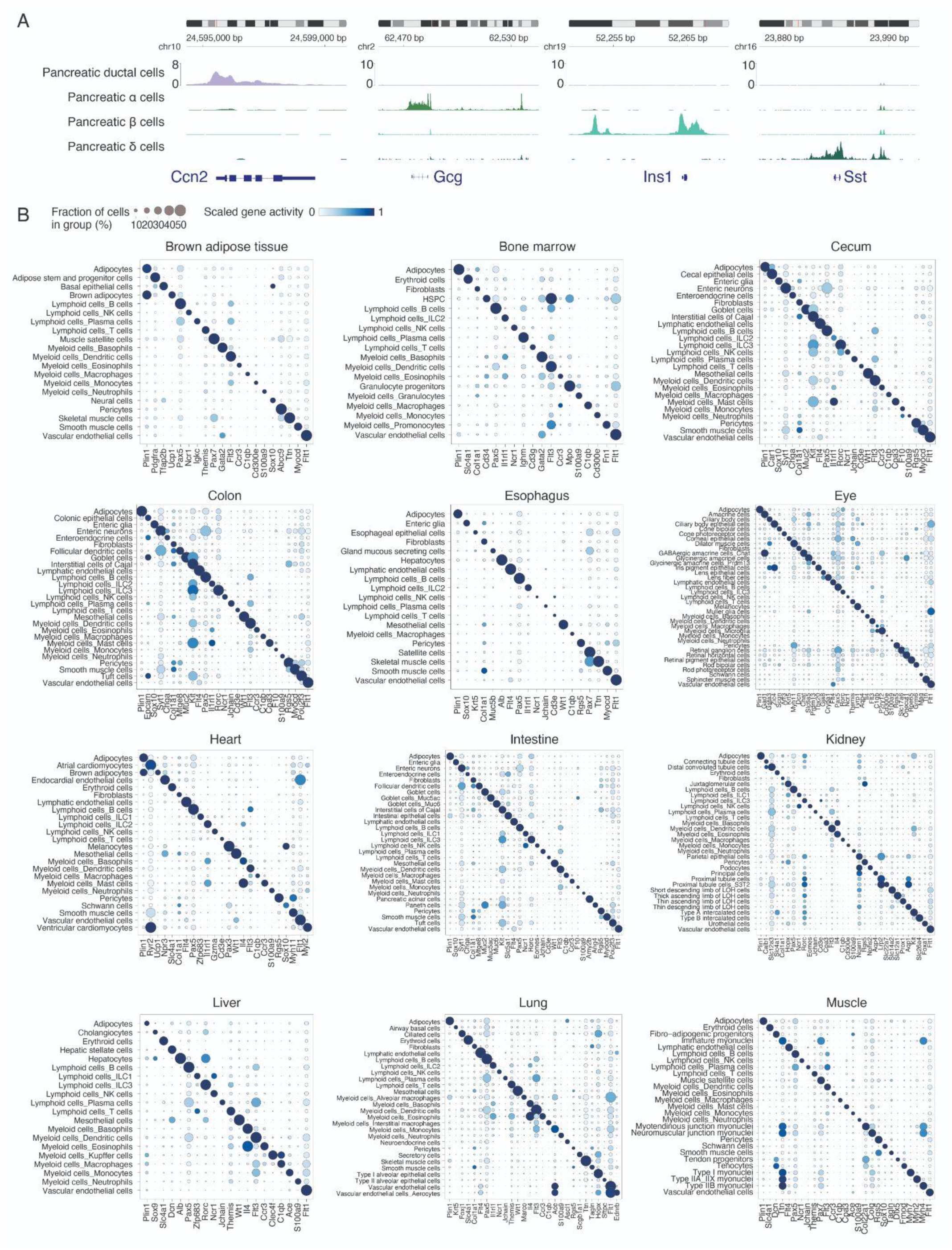
Annotation of main cell types using gene accessibilities. **(A)** Genomic tracks showing examples of cell type annotation using accessibilities of marker genes for pancreatic ductal cells (*Ccn2*), alpha cells (*Gcg*), beta cells (*Ins1*), and delta cells (*Sst*). (B) Dot plots showing gene markers used for annotating main cell types across tissues. The size of the dot encodes the percentage of cells within a cell type in which that marker was detected, and its color encodes the average accessibility level. A complete list of markers can be found in Table S2.

**Figure S3.**
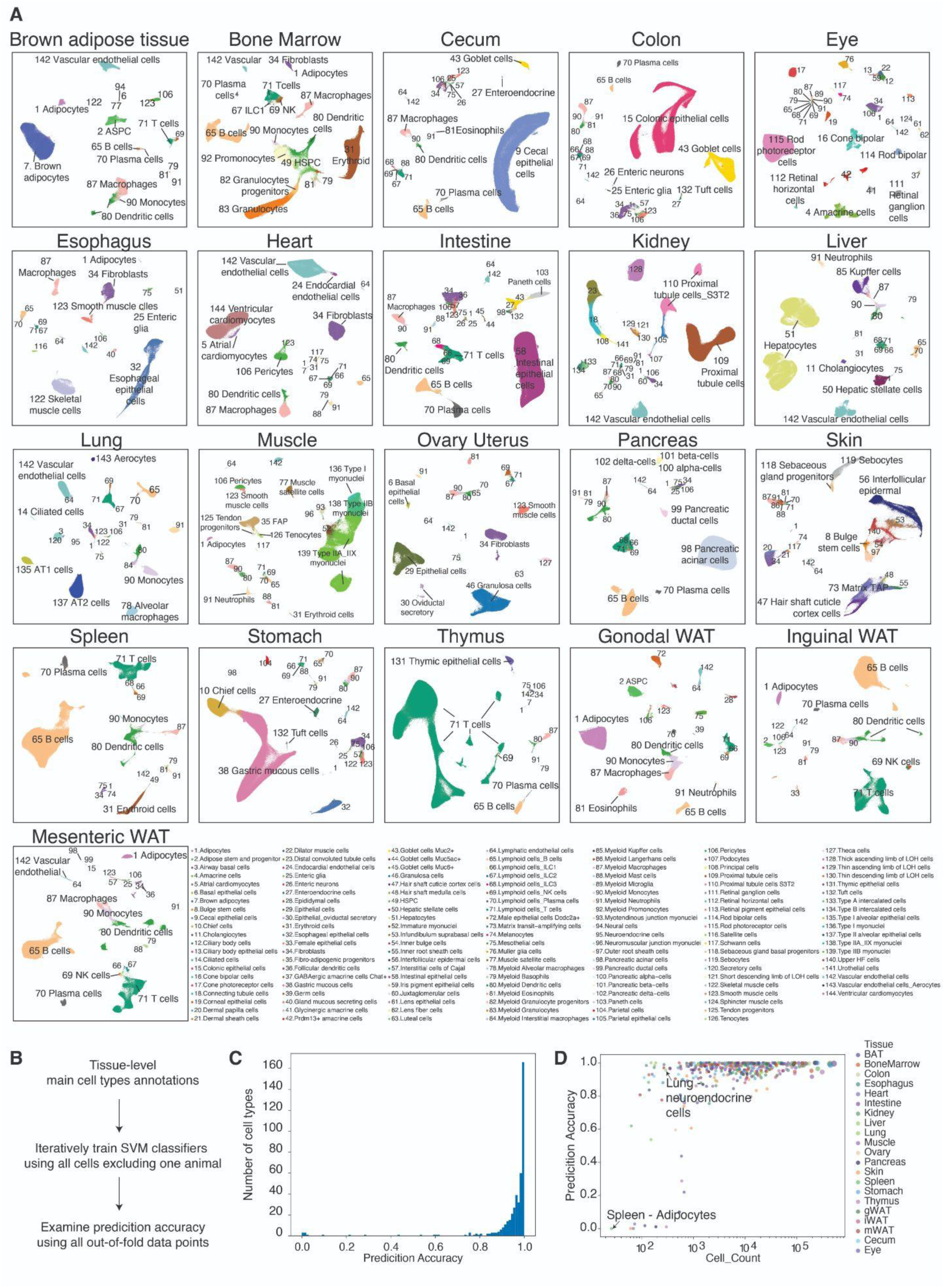
Cell type annotations and reproducibility analysis. (A) UMAP visualization of cells from each tissue, colored by unique cell types collapsed from all tissues. (B) Leave-one-animal-out cross-validation analysis workflow. For each tissue, we created a training set by taking cells from all except one animal, built SVM classifiers to predict the cell type labels from the dimension reduction matrices, and used the models to predict the labels for those cells from the hold-out animal. Such procedures were repeatedly done over all the biological replicates, and the reproducibility of cell type identification was evaluated based on the prediction accuracy of all out-of-fold data points. (C) Histogram of prediction accuracies across all tissue-level main cell types, derived from SVM classifiers trained on all but one animal and tested on the held-out replicate. (D) Scatter plot of prediction accuracy versus cell count for each cell type. Points are colored by tissue of origin.

**Figure S4.**
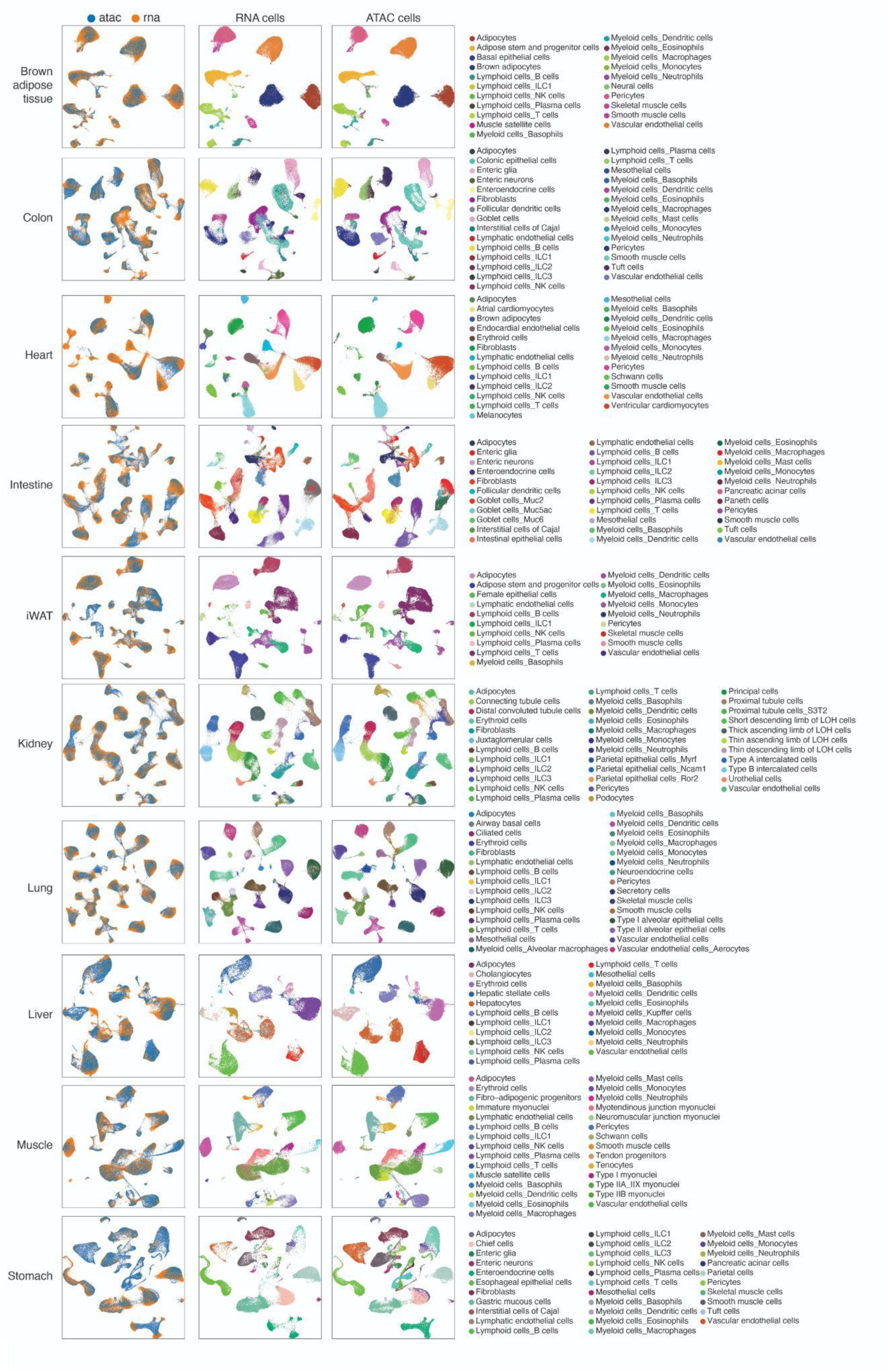
Cross-validation of ATAC-seq cell clustering and annotation using RNA-seq atlas. UMAP visualizations showing integrated snATAC-seq and snRNA-seq data across ten representative tissues, colored by molecular layers (left), RNA-seq cell annotations (middle), and ATAC-seq cell annotations (right). For each tissue, we subsampled 10,000 cells per main cell type per modality and performed integration using the standard Seurat workflow (**Methods**).

**Figure S5.**
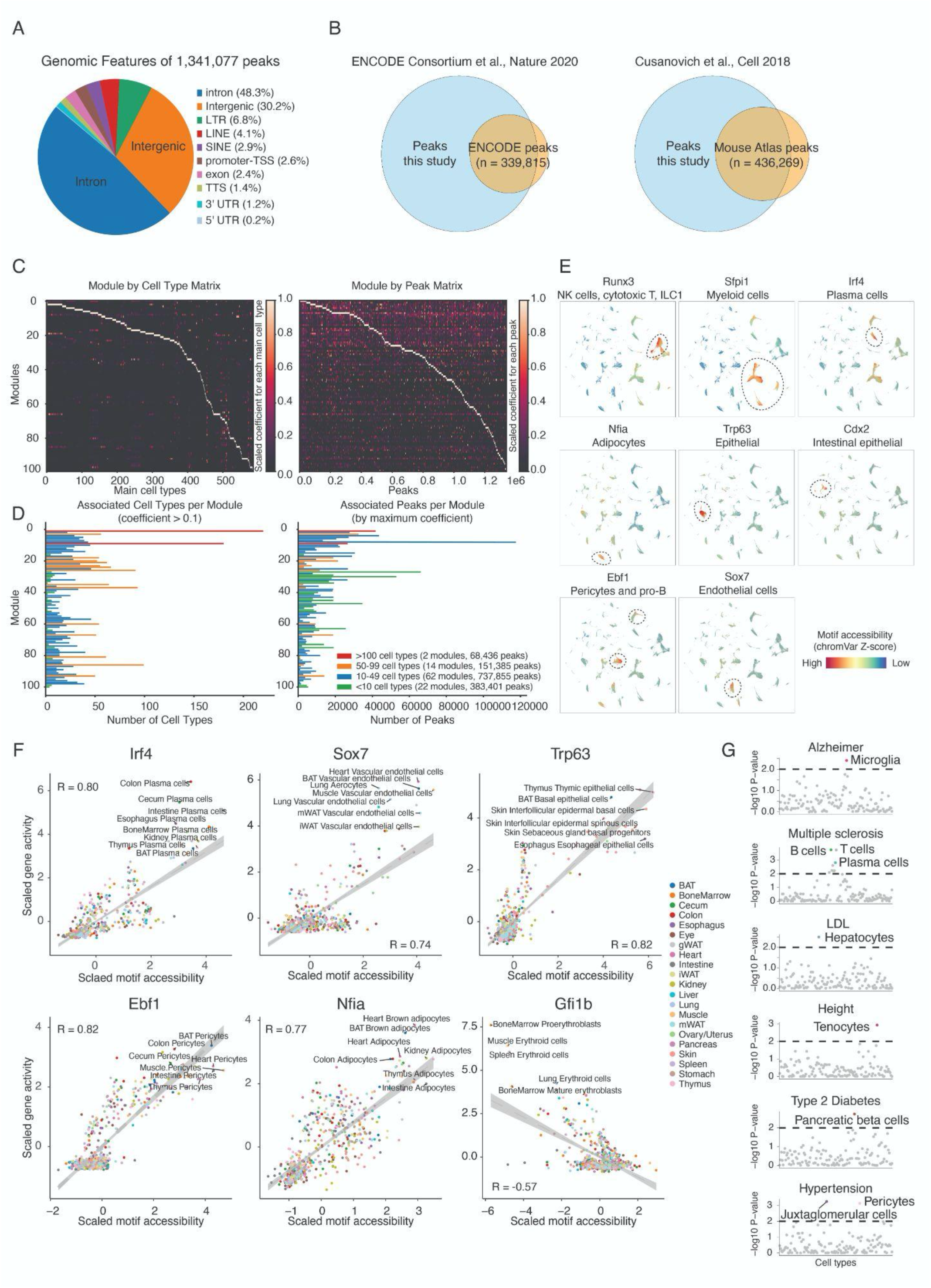
Identifications and characterizations of cell-type-specific cis-regulatory elements (CREs). **(A)** Genomic features of 1,341,077 peaks across the mouse genome. Peaks were annotated using HOMER. TSS, transcription start site; TTS, transcription termination site; UTR, untranslated region. (B) Venn plot showing the overlap between the peak set determined in this study and from the mouse ENCODE registry (*17*) and the Cusanovich et al. atlas (*12*). **(C)** Non-negative matrix factorization decomposition of the aggregated main cell type by peak matrix into 100 modules. Left: module–cell type coefficient matrix, where coefficients are scaled for each cell type. Right: module–peak coefficient matrix, with coefficients scaled for each peak. **(D)** Summary of module associations. Left: the number of cell types associated with each module, defined by coefficients >0.1 (95th percentile of the module–cell type matrix). Right: the number of peaks assigned to each module by their maximum coefficient. Modules are grouped and color-coded by the number of associated cell types. (E) UMAP plots as in Fig.1C, colored by motif accessibilities (quantified by chromVar (*19*)) of the lineage-specific transcription factor. **(F)** Scatter plots showing the example TFs whose gene accessibility levels are positively or negatively correlated with motif accessibility across cell types and tissues. Each point indicates a cell type from a specific tissue. Gene accessibilities were quantified as counts per million, and motif activities were quantified by chromVar (*19*). **(G)** Scatter plots showing the enrichments of the phenotype-associated SNPs in cell-type-specific peaks. X-axis: all unique cell types collapsed across tissues; y-axis: significance of enrichments (-log10 p-value). The top enriched cell types were labeled for each phenotype.

**Figure S6.**
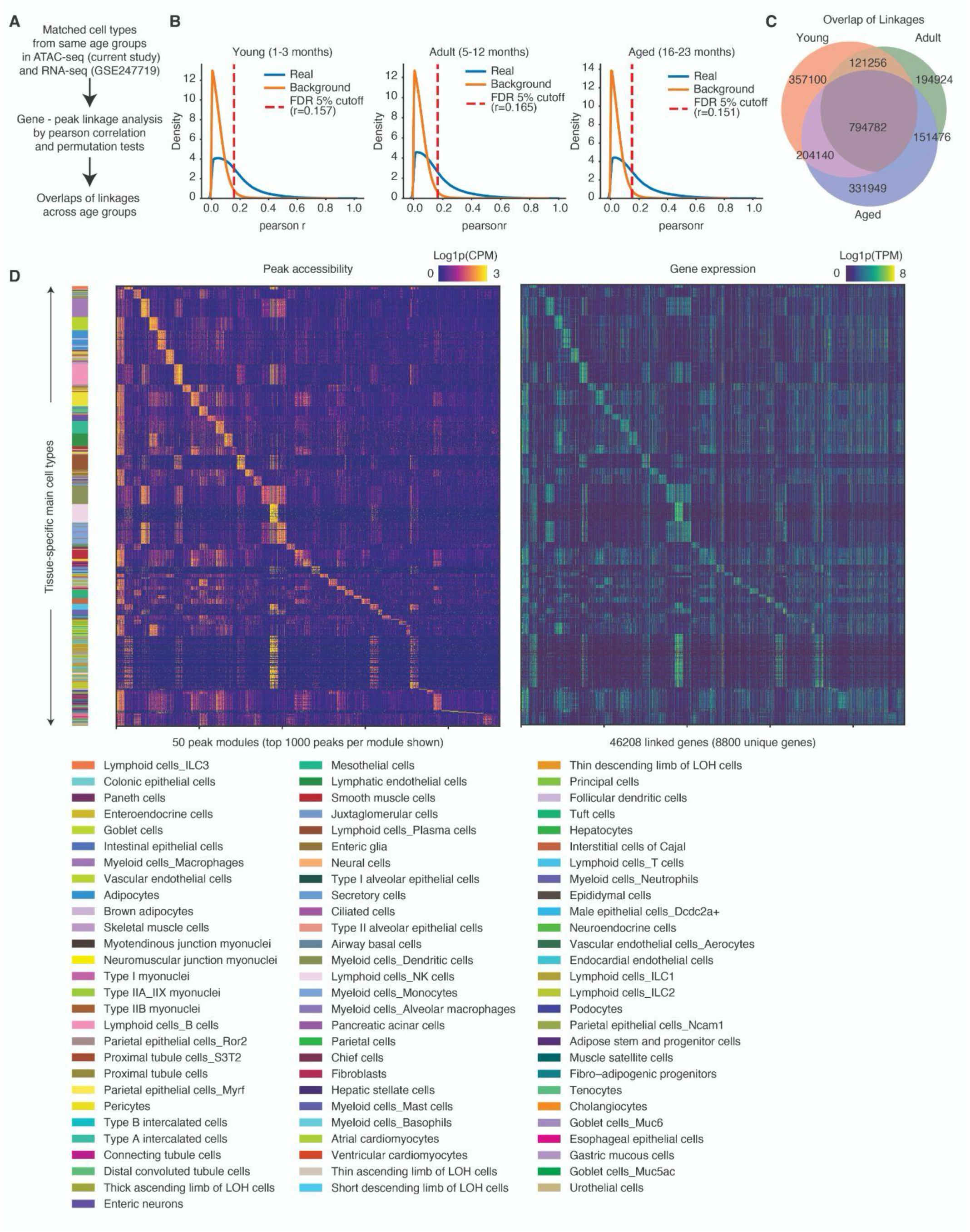
Age-stratified RNA–ATAC gene–peak linkage analysis across cell types. **(A)** Schematic of the analysis strategy. For each age group (young, adult, aged), we collapsed single-cell ATAC-seq and RNA-seq profiles by tissue and cell type to generate matched pseudo-bulk matrices. Gene–peak linkages were defined as significant Pearson correlations (within ±100 kb of the TSS) across cell types. Significance was determined against a shuffled background distribution with an FDR cutoff at 0.05. (B) Distributions of observed Pearson correlations (blue) versus background (orange) for young (1–3 months), adult (5–12 months), and aged (16–23 months) groups. The red dashed line marks the FDR cutoff. Numbers indicate thresholds used to define significant correlations. (C) Venn diagram of positively correlated gene–peak pairs identified in each age group and their overlaps. **(D)** Non-negative matrix factorization of gene–peak linkage matrices identifies 50 peak modules with cell-type-specific profiles. Heatmaps show cell-type by module activity for peak accessibility (left, log1p CPM) and expression of linked genes (right, log1p TPM). Cell types are grouped vertically, with color bars denoting cell type identities.

**Figure S7.**
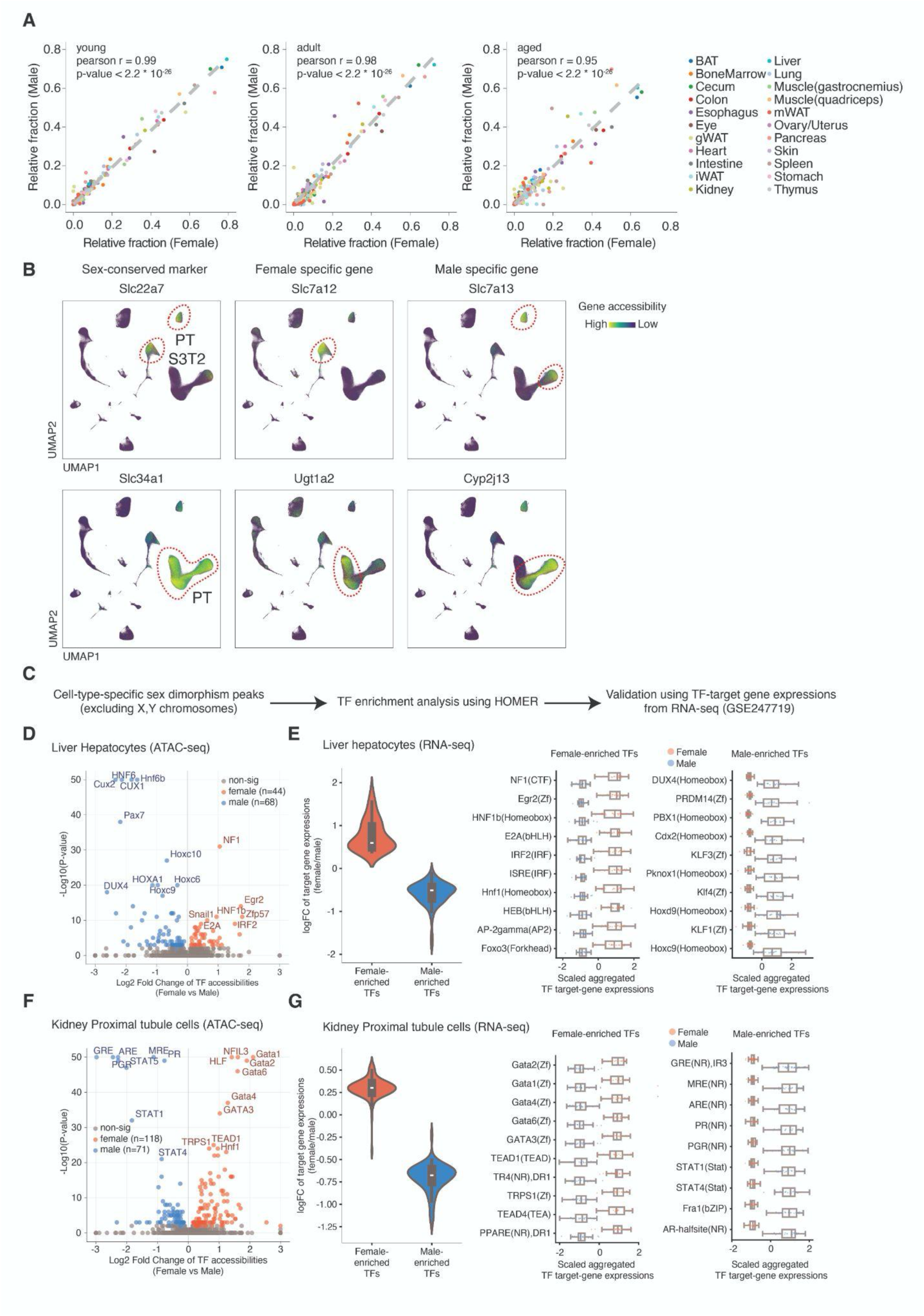
Examinations of sex dimorphism in cell proportion and molecular states. (A) Scatterplot showing the fraction of each main cell type in each tissue between males and females, stratified by age groups. Each dot represents a cell type in a specific tissue. (B) UMAP plots of all kidney cells, colored by accessibility of genes shared between sexes (left; *Slc34a1* for general proximal tubule cells (PT), *Slc22a7* for PT S3T2), unique to females (middle) and males (right). (C) Analysis workflow to identify sex-dimorphic transcription factors. Sex-biased chromatin accessibility peaks (excluding sex chromosomes) were identified across all ages, followed by TF motif enrichment analysis using HOMER and validation using RNA-seq–based TF target gene expression. (D) Volcano plot of TF motif enrichments from sex-dimorphic peaks in liver hepatocytes (ATAC-seq). Blue, male-biased motifs; red, female-biased motifs; grey, non-significant. Selected TF motifs are labeled. (E) RNA-seq validation using sex-biased gene expressions in liver hepatocytes. Left, violin plots showing the distribution of target gene expression (log fold-change, female vs. male) for female- vs. male-enriched TFs. Right, boxplots of aggregated target gene expression values for representative TFs. (F) Volcano plot of TF motif enrichments from sex-dimorphic peaks in kidney proximal tubule cells (ATAC-seq). Blue, male-biased motifs; red, female-biased motifs; grey, non-significant. Selected TF motifs are labeled. (G) RNA-seq validation in kidney proximal tubule cells. Left, violin plots of TF target gene expression as in E. Right, boxplots of aggregated target gene expression values for representative TFs.

**Figure S8.**
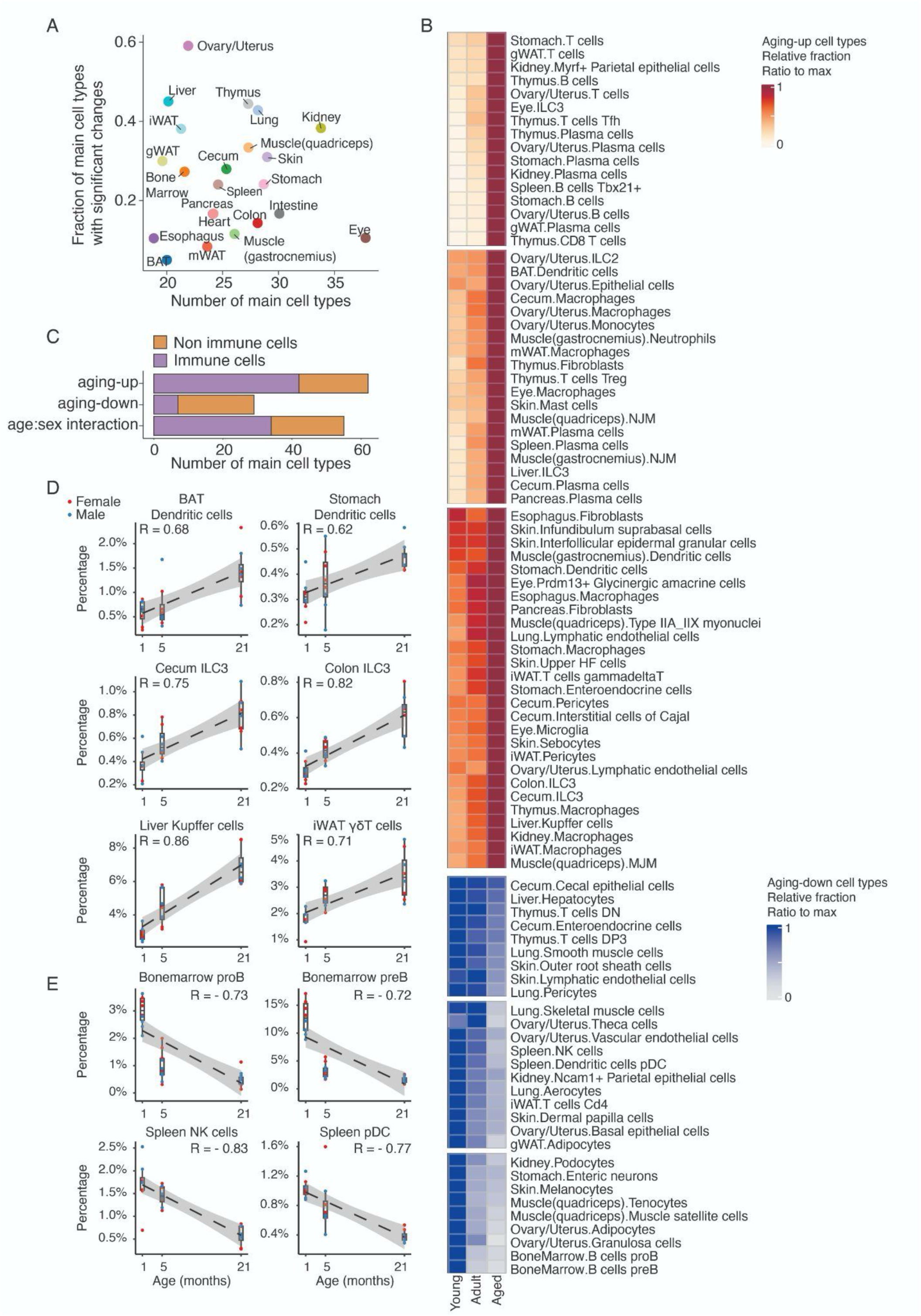
Aging-associated changes in main cell type proportions across tissues. (A) Scatterplot showing the fraction of main cell types whose proportions changed significantly with age or age-sex interaction in each tissue. **(B)** Heatmaps visualizing proportional changes in all aging-associated, sex-independent main cell types. The fraction of each main cell type within its tissue of origin was calculated per sample, averaged within each age group, and normalized to the most abundant condition. Cell types were ordered by hierarchical clustering implemented in ComplexHeatmap (*87*). (C) Bar plot showing the number of main cell types derived from immune or non-immune cell types within aging-associated, sex-independent (aging-up and aging-down), or age-sex interaction groups. (D-E) Scatter and box plots showing examples of immune cell types that expand (D) or decline (E) with age, with a linear regression line (and a Pearson correlation coefficient). Each dot represents the cell-type specific proportion of the cell type in each animal.

**Figure S9.**
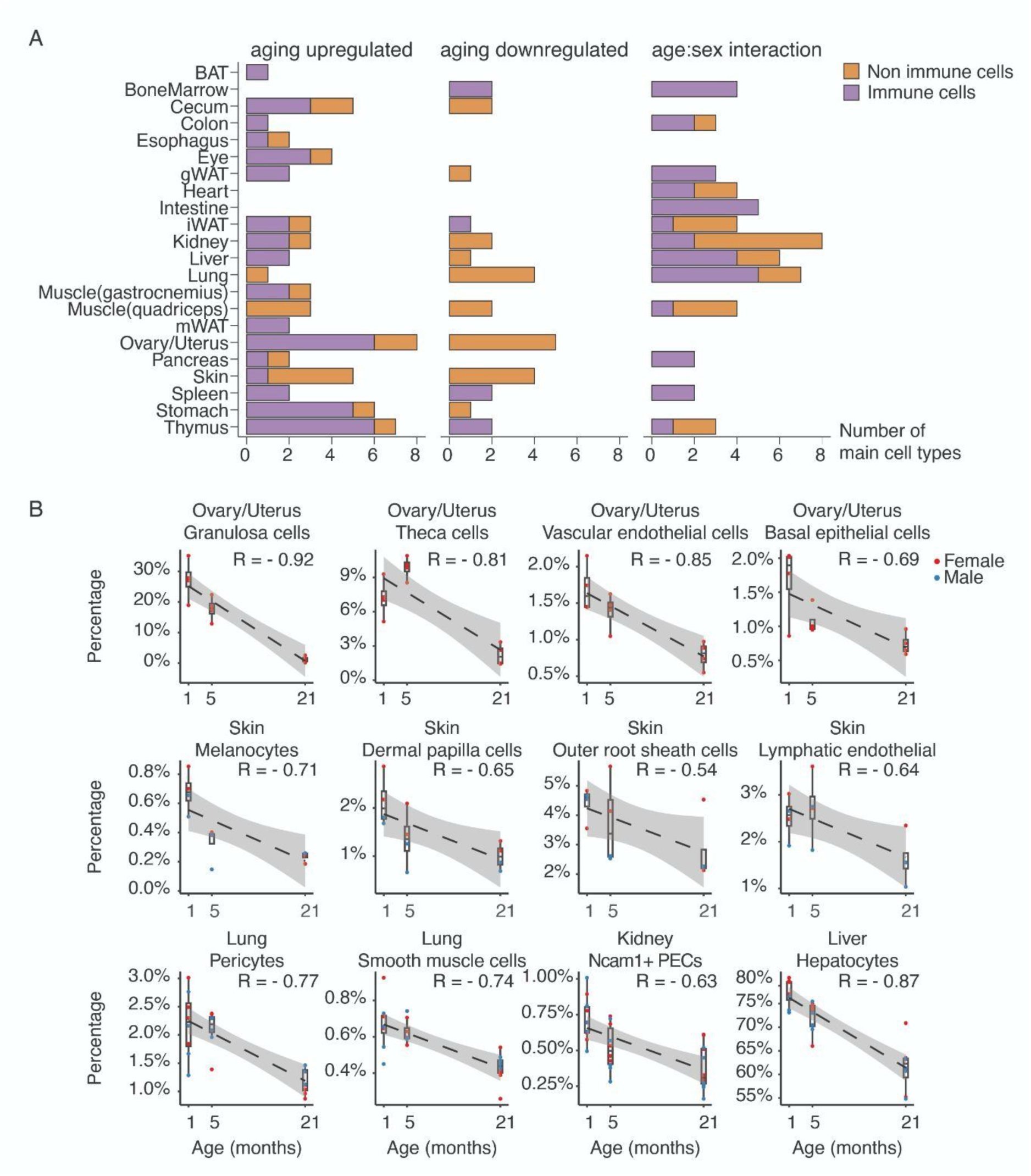
Aging-associated depletion of functional cell types across tissues. (A) Bar plot showing the number of main cell types derived from immune cell types or non-immune cell types within aging-associated, sex-independent (aging-up and aging-down), or age-sex interaction groups for each tissue. (B) Scatterplot showing examples of main cell types that decline with age across tissues, with a linear regression line (and a Pearson correlation coefficient). Each dot represents the cell-type-specific proportion of the cell type in each animal.

**Figure S10.**
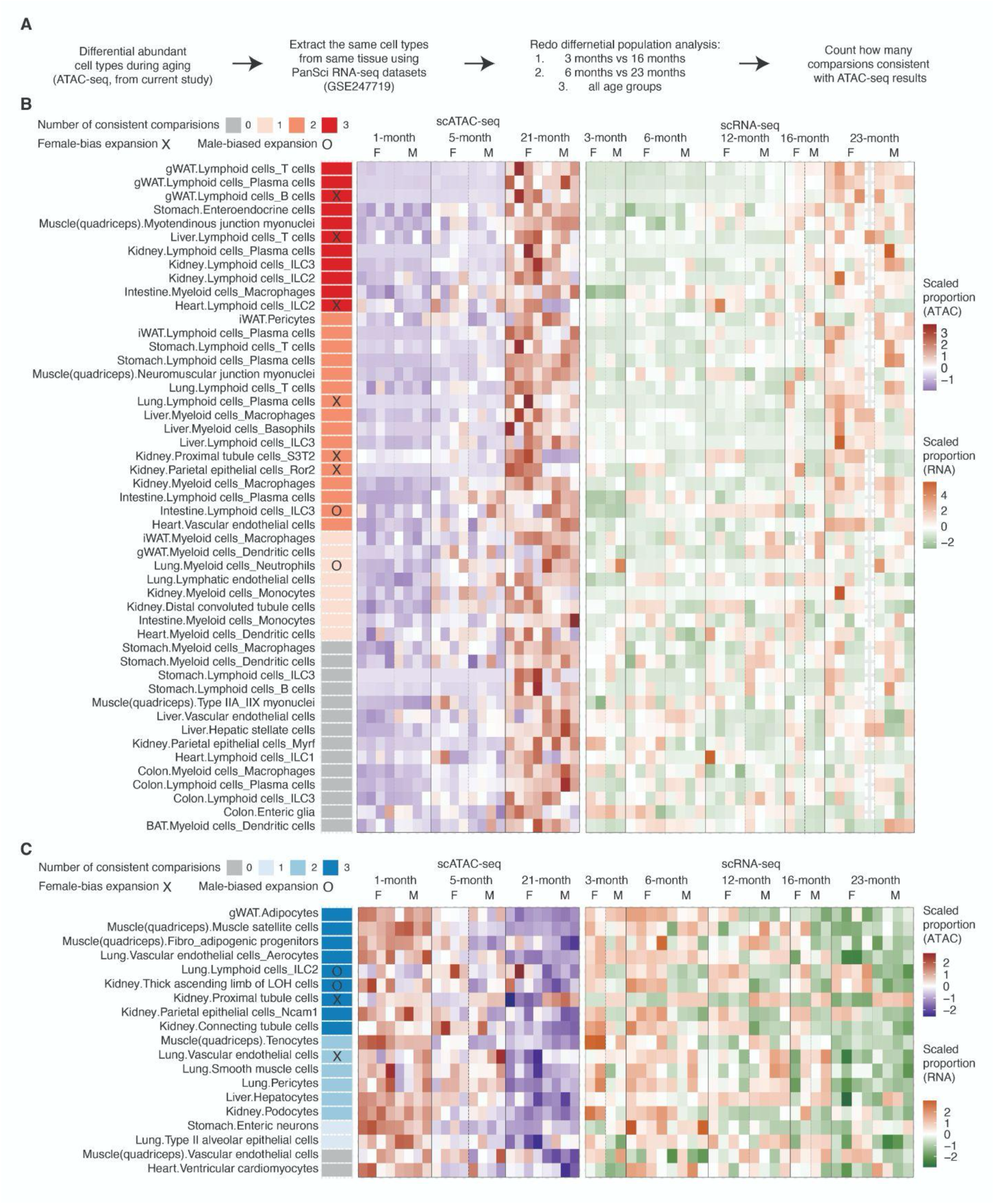
Cross-validation of age- and sex-associated cell type population changes between scATAC-seq and the PanSci scRNA-seq atlas of mouse aging. (A) Schematic of the analysis strategy. For each aging-associated cell type identified in the current study, we extracted the corresponding cells from the same tissue in PanSci RNA-seq and performed differential-abundance analyses. Because the scRNA-seq data include five ages (3, 6, 12, 16, and 23 months) collected in two experimental batches (3 and 16 months in one batch; 6, 12, and 23 months in the other), we ran three DA analyses comparing age groups to minimize potential batch effects. We considered a change replicated when at least one comparison yielded p < 0.05 with a concordant logFC sign relative to sc-ATAC-seq. For age-by-sex effects, we considered a cell type validated when the interaction coefficient was significant (p-value < 0.05) and the direction matched the sc-ATAC-seq result. (B) Heatmap showing cell types with significant increases in abundance with age in scATAC-seq, shown alongside corresponding scRNA-seq results. Rows represent cell type. Columns represent biological replicates. Left annotation indicates the number of comparisons replicated in scRNA-seq, with symbols denoting sex-biased expansions (X = female-biased, O = male-biased). (C) Heatmap showing cell types with significant decreases in abundance with age in scATAC-seq, analyzed in the same manner as (B), with annotation bars indicating the degree of replication across comparisons.

**Figure S11.**
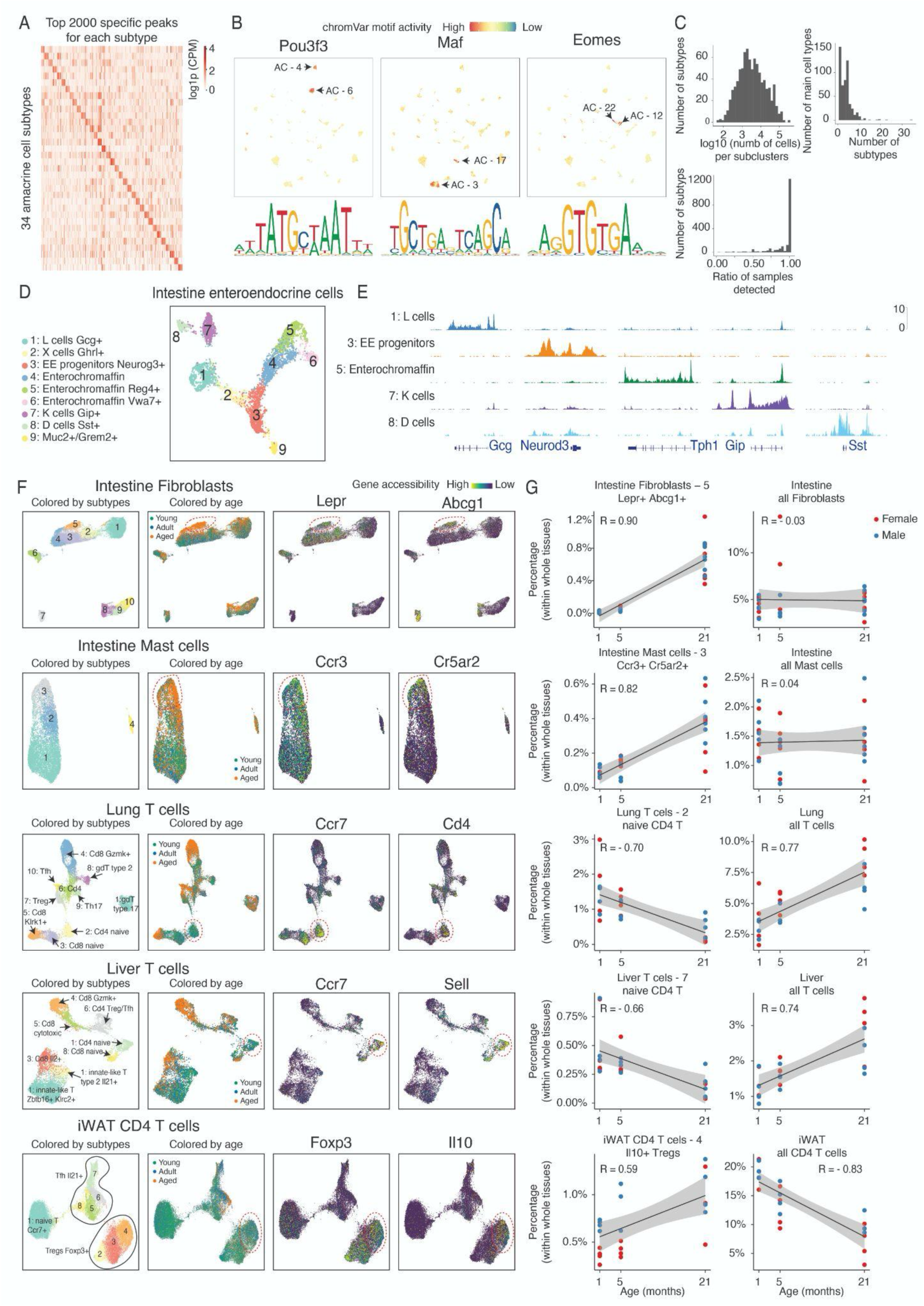
Identification of cell subtypes and associated population changes in aging. (A) Heatmap showing the aggregated accessibility of peaks specific to each main subtype in amacrine cells, quantified by counts per million. (B) UMAP visualization of eye amacrine cells, colored by motif accessibilities of example transcription factors specific to distinct subtypes. Motif activities were quantified by chromVar (*19*). (C) Histograms showing the distribution of cell numbers per subtype (Top left), the number of subtypes per main cell type (Top right), and the fraction of tissue samples that contain cells from each cell subtype (Bottom left). (D) UMAP visualization of intestine enteroendocrine cells, colored by subtype identity. (E) Genomic tracks showing the gene accessibilities marking different subtypes of intestine enteroendocrine cells. **(F)** UMAP plots showing subclustering results for intestine fibroblasts, intestine mast cells, lung T cells, liver T cells, and iWAT CD4 T cells, colored by subtype ID, age group, and accessibilities of genes marking circled subclusters. (G) Scatterplot showing the proportion changes of indicated subclusters and their parental main cell types with age, along with a linear regression line.

**Figure S12.**
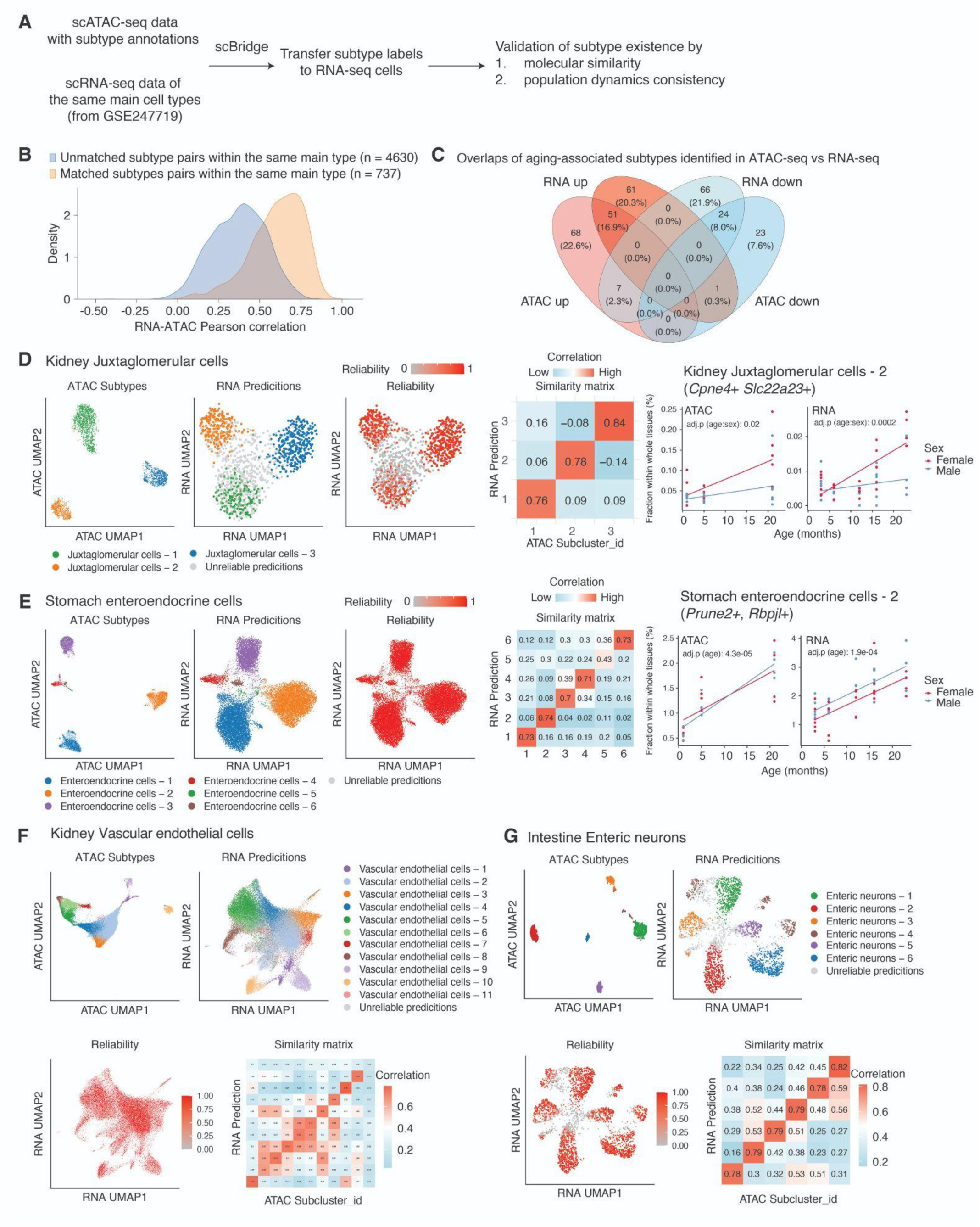
Cross-modality validation of cell subtypes identity and aging-associated changes. (A) Schematic of the analysis workflow. Subtype annotations identified in scATAC-seq were transferred to scRNA-seq cells of the same main cell types (*7*) using scBridge(*58*). Subtype existence was validated by molecular similarity and population dynamics consistency. (B) Density distribution of RNA–ATAC Pearson correlations for unmatched versus matched subtype pairs within the same main cell type. Matched pairs (orange) show higher correlations compared to unmatched pairs (blue). (C) Venn diagrams showing overlaps of aging-associated subtypes identified in ATAC-seq versus RNA-seq. **(D–G)** Representative examples of subtypes validated across modalities: kidney juxtaglomerular cells (D), stomach enteroendocrine cells (E), vascular endothelial cells (F), and intestinal enteric neurons (G). For each case, UMAP visualizations of ATAC subtypes, RNA predictions, and prediction reliability are shown, along with subtype similarity matrices. For (D) and (E), line plots show consistent age-associated compositional changes across ATAC-seq and RNA-seq data, stratified by sex. Each dot represents one biological replicate.

**Figure S13.**
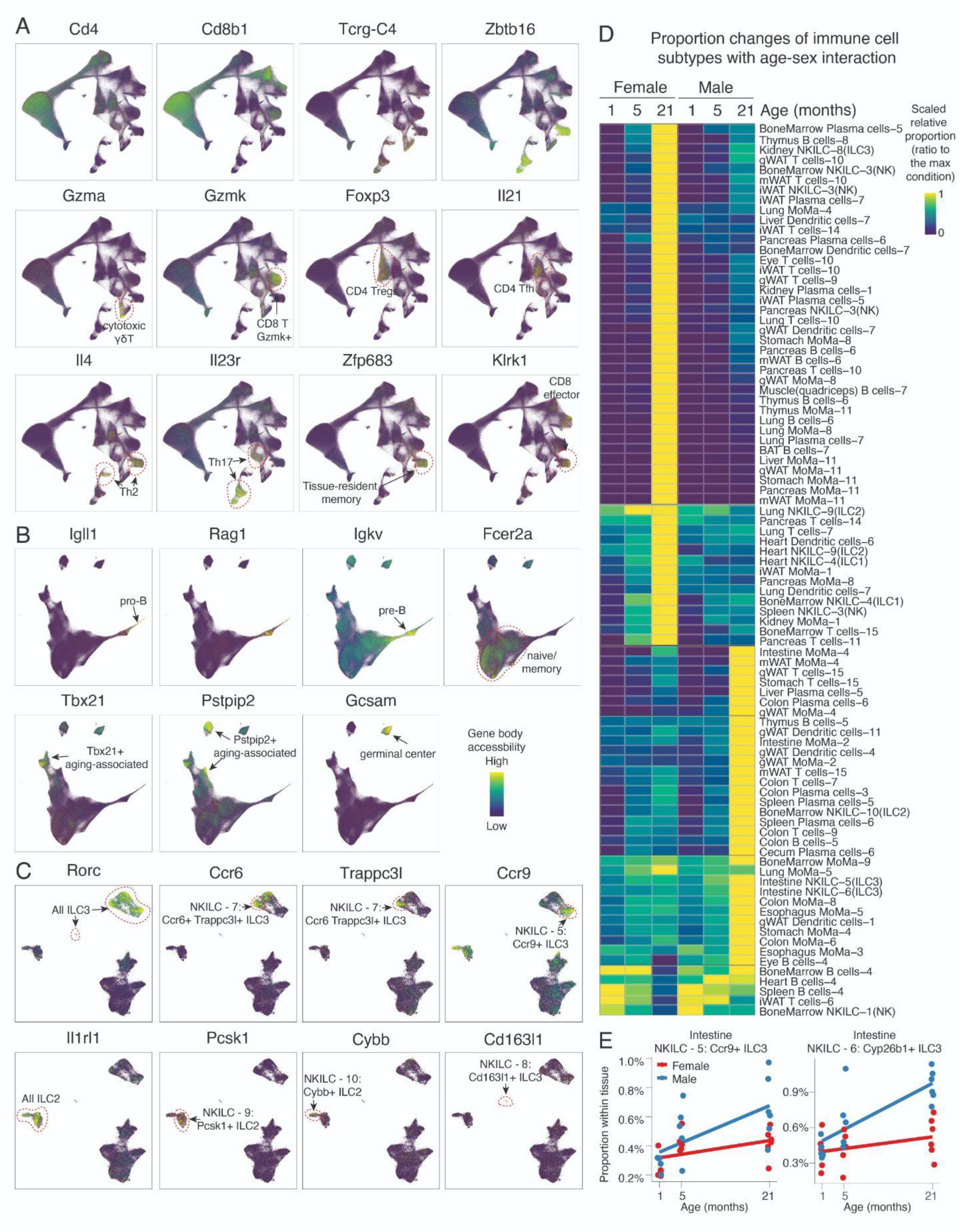
Global characterizations of immune cell states and their proportional changes with age. (A) UMAP plots showing combined clustering of all T cells across tissues, colored by accessibilities of genes marking distinct subtypes. (B) UMAP plots showing combined clustering of all B cells across tissues, colored by accessibilities of genes marking distinct subtypes. (C) UMAP plots showing combined clustering of all innate lymphoid cells across tissues, including NK cells, ILC1, ILC2 and ILC3, colored by accessibilities of genes marking distinct subtypes. (D) Heatmap of relative proportions of immune cell subtypes with significant age-sex interactions. Rows represent immune cell subtypes within their respective tissues, while columns correspond to mouse conditions grouped by age and sex. For each immune cell subtype within each tissue, its proportion (within each tissue) were first averaged across samples and then normalized to the maximum value within each row. (E) Scatterplot showing male-baised expansions of *Ccr9*+ ILC3 and *Ccr6*+ *Cyp26b1*+ ILC3 in intestine along aging. Each dot represents one animal, with linear regression lines added for each sex.

**Figure S14.**
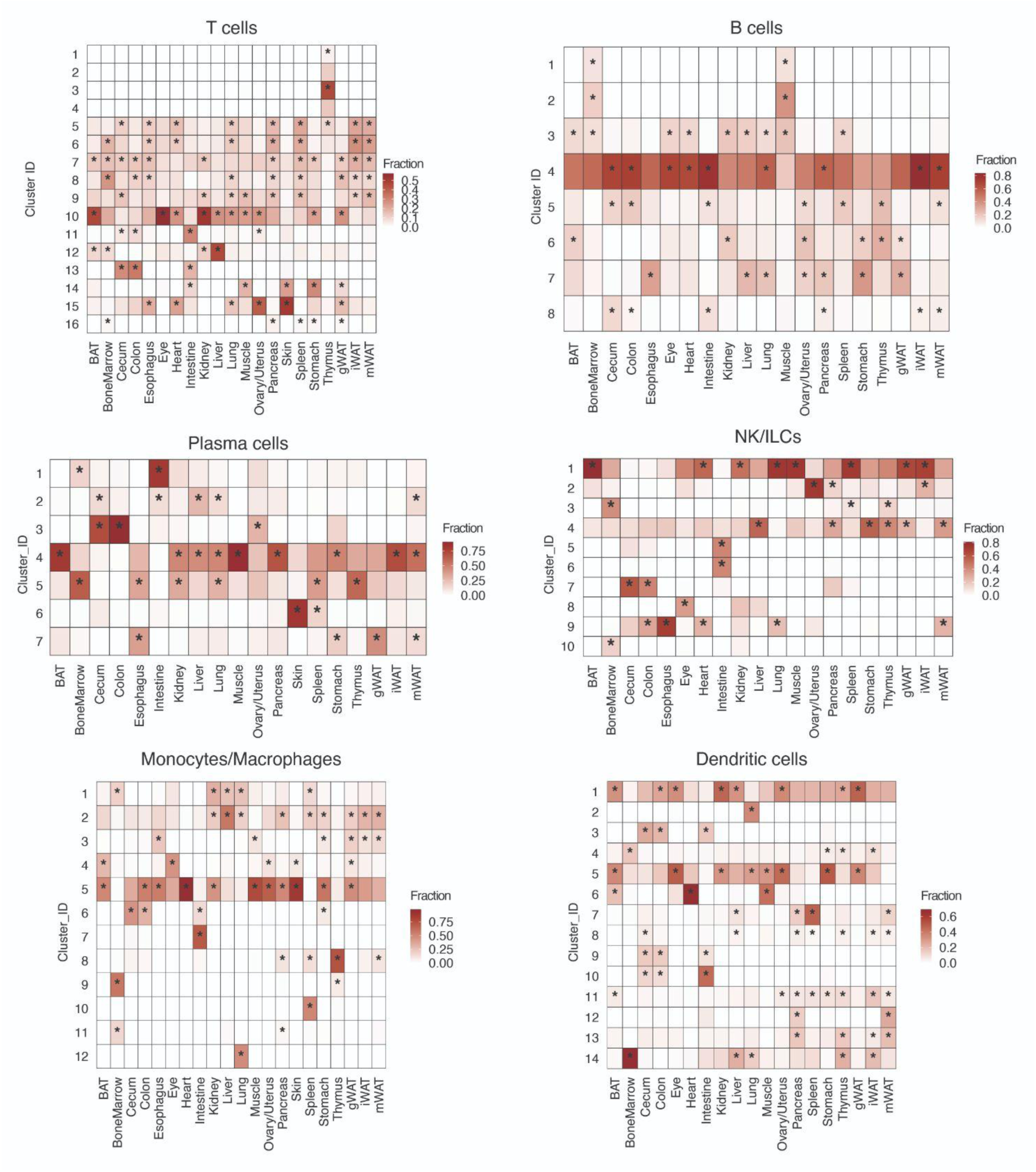
Relative proportions of immune subtypes across organs. Heatmaps show the relative fraction of each immune cell subtype within each tissue. For each subtype, asterisks mark significantly higher fractions (FDR<0.05, one-sided binomial test) in a given tissue relative to the average fraction across tissues.

**Figure S15.**
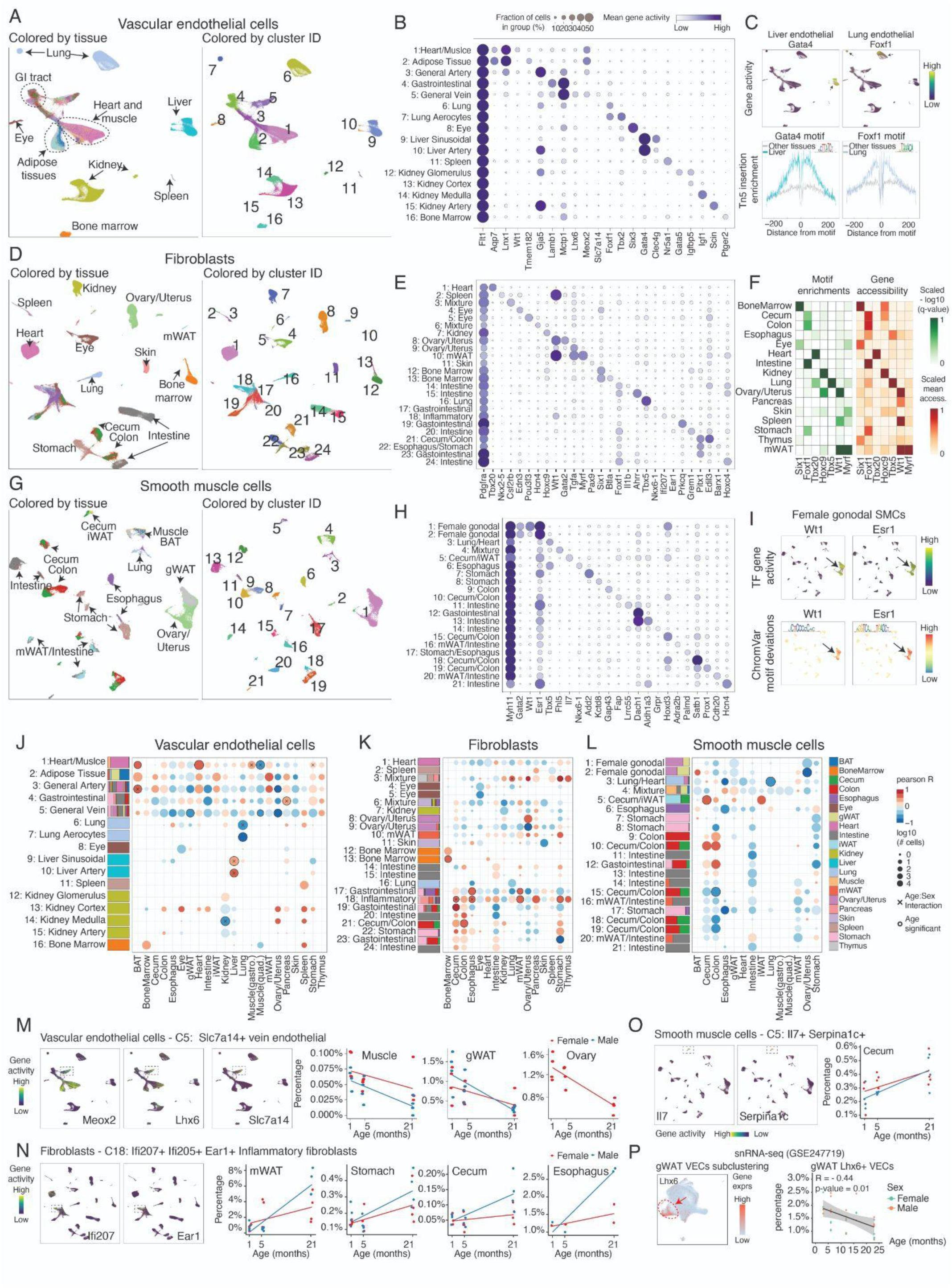
Aging-associated dynamics of vascular endothelial cells, fibroblasts, and smooth muscle cells across organs. (A) UMAP visualization showing the combined clustering for vascular endothelial cells (VECs, top), fibroblasts (FBs, middle) and smooth muscle cells (SMCs, bottom) across tissues, colored by tissue origin (Left) and cluster ID (Right). (B) Dot plots showing the accessibility of marker genes for cell clusters of VEC (top), FB (middle) and SMC (bottom), including a general marker for each cell type (*Flt1* for VEC, *Pdgfra* for FBs, *Myh11* for SMCs) (C) UMAP plot of VECs colored by accessibilities of the liver VEC marker *Gata4* and lung VEC marker *Foxf1* (D) Motif footprinting analysis showing enrichment of GATA4 motifs in liver VECs and FOXF1 motifs in lung VECs compared with other tissues. (E) Heatmaps showing the motif enrichments and gene body accessibility of tissue-specific transcription factors. Motif enrichment scores (-log10 q-value) were computed with HOMER (*77*) using tissue-specific peaks and scaled to the top enriched tissue. (F) UMAP plots of SMCs colored by gene accessibility (Top) and chromVar motif deviation scores (Bottom) for gonadal SMC markers *Wt1* and *Esr1*. (G) The stacking bar plots show tissue composition per cluster for VECs (Left), FBs (Middle), and SMCs (Right). The dot plots show aging-associated proportional changes of clusters across tissues, colored by Pearson correlation between cluster proportion (normalized per tissue) and age. Clusters with significant age/age-sex effects are annotated (p-adjusted < 0.05). (H) UMAP of VECs colored by the accessibility of VEC-5 markers (*Meox2*, *Lhx6*, *Slc7a14*). (I) Scatterplots showing the age-related decline in VEC-5 proportions in muscle (quadriceps), gonadal white adipose tissue (gWAT), and ovary/uterus. (J) UMAP of FBs colored by the accessibility of FB-18 markers (*Ifi207*, *Ear1*). **(K)** Scatterplots showing the age-associated increase in FB-5 proportions in mesenteric WAT (mWAT), stomach, cecum, and esophagus. (L) UMAP plot showing gWAT VEC cells isolated from snRNA-seq data (*7*), colored by *Lhx6* expression (VEC-5 marker). (M) Scatterplot showing age-dependent reduction in *Lhx6*+ gWAT endothelial cells.

**Figure S16.**
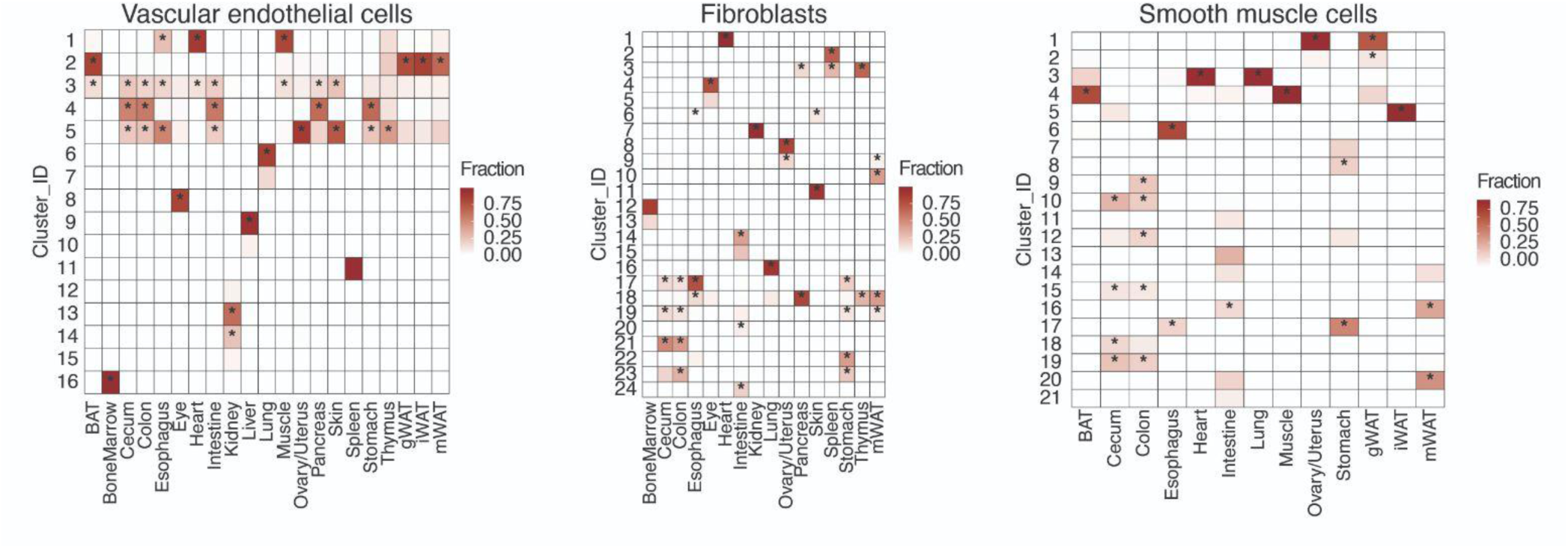
Relative proportions of endothelial cells, fibroblasts and smooth muscle subtypes across organs. Heatmaps show the relative fraction of each immune cell subtypes within each tissue. For each subtype, asterisks mark significantly higher fractions (FDR<0.05, one-side binomial test) in a given tissue relative to the average fraction across tissues.

**Figure S17.**
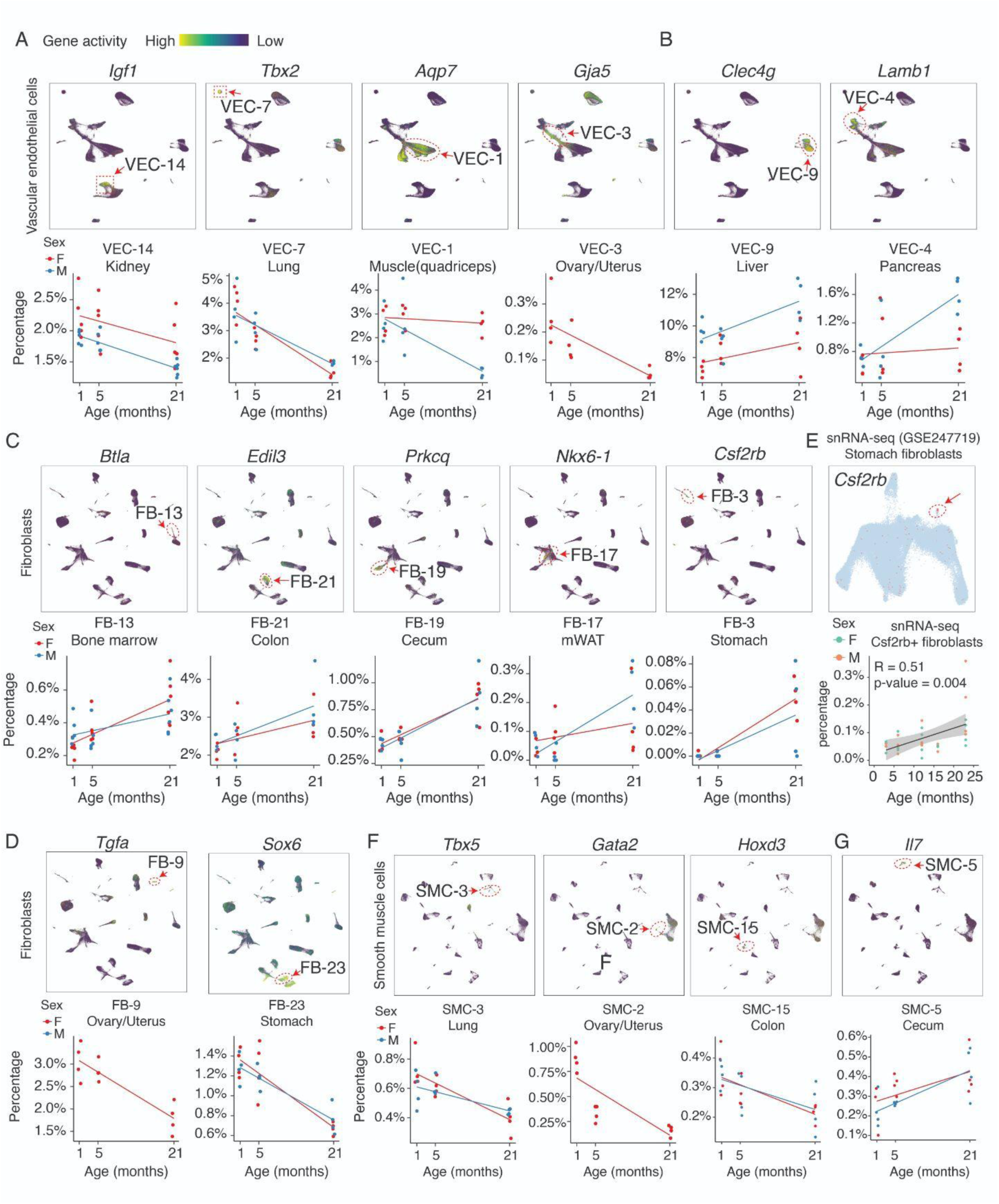
Cross-tissue identifications of subtypes for vascular endothelial cells, fibroblasts, smooth muscle cells, and their aging-associated population changes. (A-B) Top: UMAP plots showing the combined clustering results of vascular endothelial cells, colored by the accessibility of genes representing subtypes that expand (A) or deplete (B) with age. Bottom: Scatterplots showing the proportions of the indicated subtypes (normalized to total cells within the tissue) across three age groups, with linear regression lines added for each sex. (C-D) UMAP plots showing the combined clustering results of fibroblasts, colored by the accessibility of genes representing subtypes that expand (C) or deplete (D) with age. Bottom: Scatterplots showing the proportions of the indicated subtypes (normalized to total cells within the tissue) across three age groups, with linear regression lines added for each sex. **(E)** Validation of *Csf2rb*+ fibroblast expansion, as shown in (C), in aged stomach tissue using published snRNA-seq data (*7*). Left: UMAP plot of stomach fibroblasts colored by *Csf2rb* expression. Right: Scatterplot showing age-dependent expansion in *Csf2rb*+ fibroblasts, together with a linear regression line. **(F-G)** Top: UMAP plots showing the combined clustering results of smooth muscle cells, colored by the accessibility of genes representing subtypes that deplete (F) or expand (G) with age. Bottom: Scatterplots showing the proportions of the indicated subtypes (normalized to total cells within the tissue) across three age groups, with linear regression lines added for each sex.

**Figure S18.**
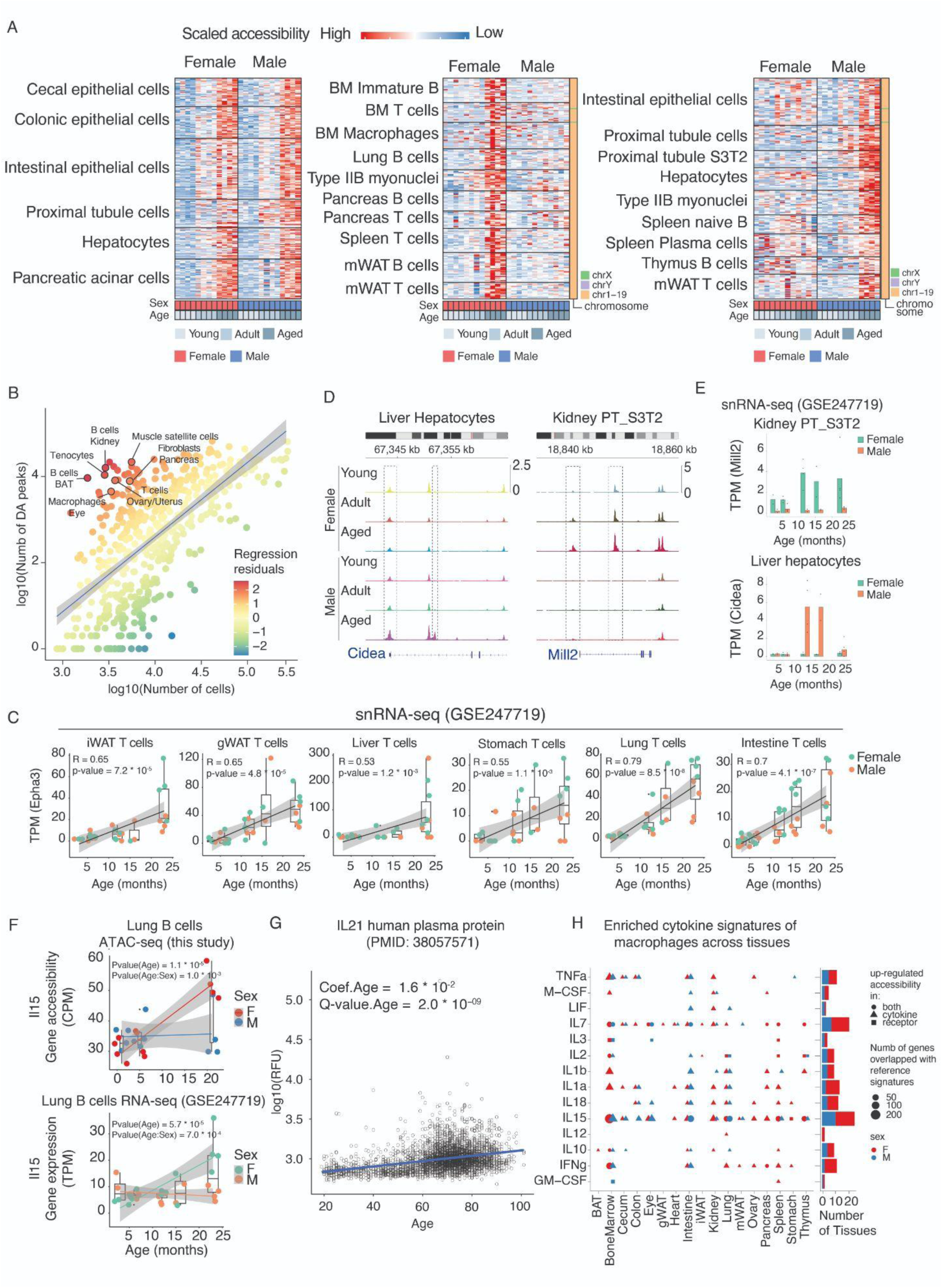
Identification of cell-type-specific molecular changes in aging. (A) Heatmaps showing examples of differentially accessible (DA) peaks that are sex-shared (left), female-specific (middle), and male-specific (right). Rows correspond to DA peaks identified in each main cell type, while columns represent individual mice. Peak accessibility was quantified as counts per million for each sample in each cell type. **(B)** Scatterplot showing the correlation between the number of cells and the number of DA peaks across cell types, with a linear regression fit. Cell types with high regression residuals are labeled. (C) Scatterplot showing the increased expression of *Epha3* in T cells across multiple tissues, with a linear regression line, validating the chromatin changes in Figure 5F. (D) Genomic tracks of male-specific DA peak in liver hepatocytes (left) and female-specific DA peak in kidney proximal tubule cells S3T2 (right). (E) Barplot showing validation of sex-specific chromatin accessibility changes in (D) using gene expression data of the same cell type. The dot represents animals. (F) Scatterplots showing the increased gene accessibility (top) and expression (bottom) of *Il15* in lung B cells, specifically in females, with linear regression lines added for each sex. **(G)** Scatterplot showing the age-related increase in plasma protein levels of IL-21. Data was obtained from (*3*), with a linear regression line included. (**H**) Left: Dot plot summarizing cytokine signatures enriched in aged macrophages across tissues. Only signatures supported by accessibility changes in secretion or receptor genes are shown. Right: Bar plot summarizing the number of tissues exhibiting activated signatures for each cytokine, separated by sex.

**Figure S19.**
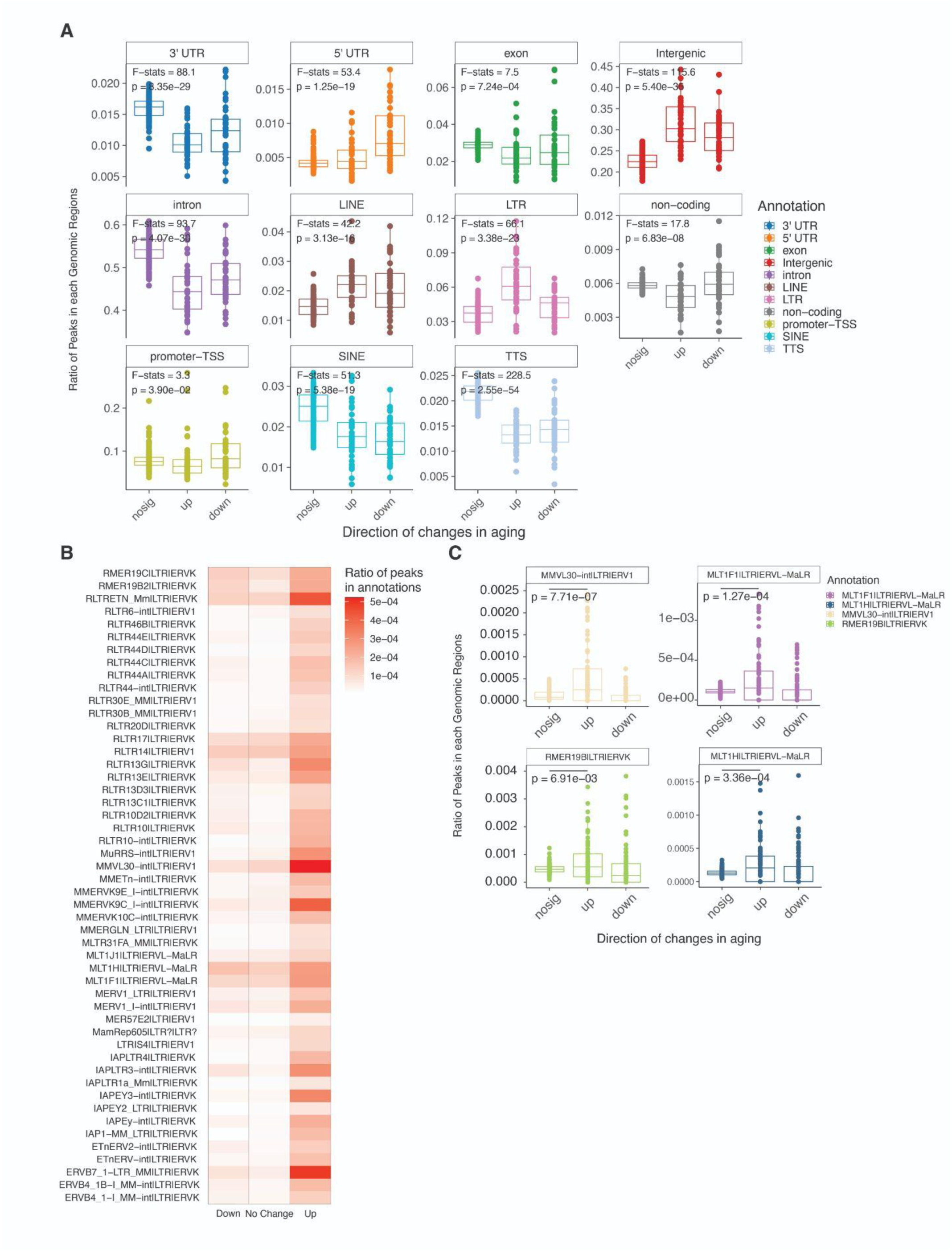
Genomic distribution of aging-associated accessibility changes. (A) Boxplots showing the proportion of peaks mapping to different genomic categories among non-changing, up-regulated, and down-regulated peaks along aging. **(B)** Heatmaps showing the mean proportions of LTR peaks exhibited significant enrichment in aging-up peaks (p-value < 0.05, logFC > 0, t-tests). (C) Boxplots showing detailed analysis of specific LTR subfamilies exhibiting significant enrichment in age-increased peaks compared to non-changing peaks.

**Figure S20.**
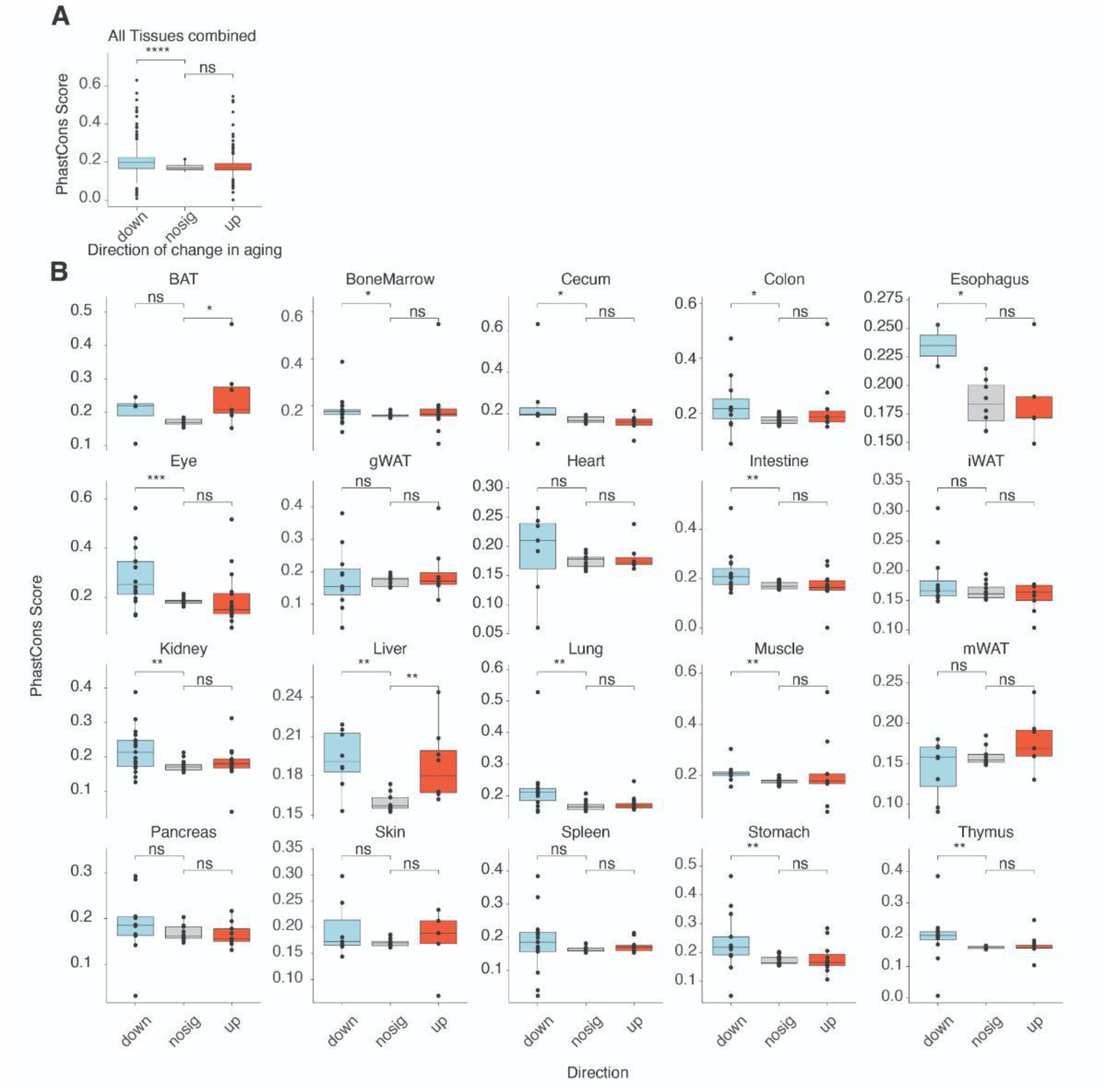
Evolutionary conservation of aging-associated accessible regions. **(A)** Boxplots showing average conservation scores for sex-shared differentially accessible (DA) peaks aggregated across all tissues. Aging-decreased peaks exhibited significantly higher conservation than stable peaks (p-value < 0.05, Wilcoxon rank-sum test). (B) Conservation score comparisons stratified by tissue. The pattern of higher conservation in aging-decreased peaks relative to stable peaks was observed consistently across many tissues.

**Figure S21.**
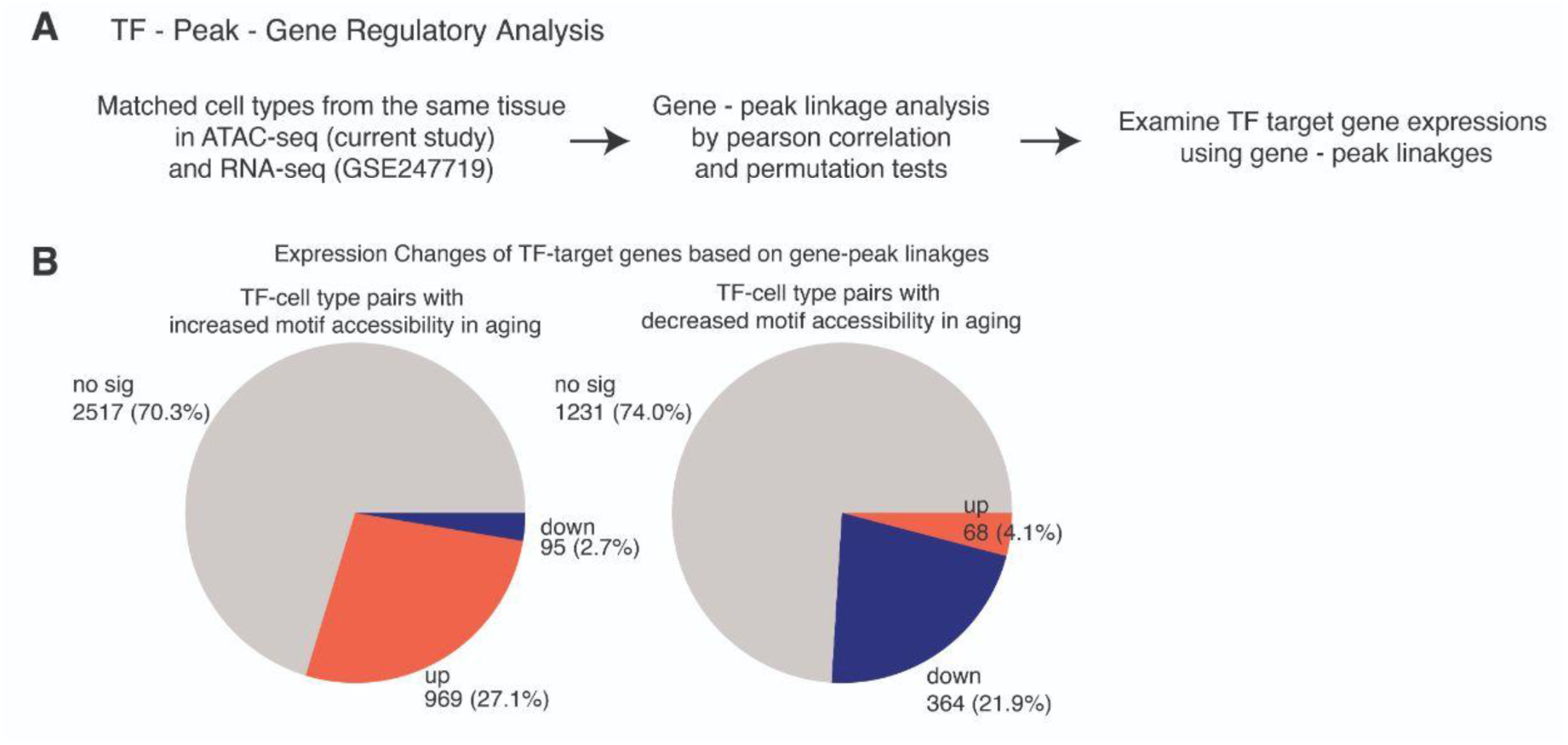
Expression changes of target genes regulated by age-influenced transcription factors. (A) Analysis workflow. For each cell type, we identified binding sites of age-influenced TFs on aging-associated peaks and used gene-peak linkages (Fig. S6) to define each TF’s target genes. We then computed pseudo-bulk expression per animal from the PanSci scRNA-seq dataset, and compared the aggregated normalized expression of TF target genes between 6- vs. 23-month-old mice using Wilcoxon rank-sum tests. (B) Pie charts show the distribution of expression changes for TF–cell type pairs with increased (left) or decreased (right) motif accessibility in aging (6 vs. 23 months).

**Figure S22.**
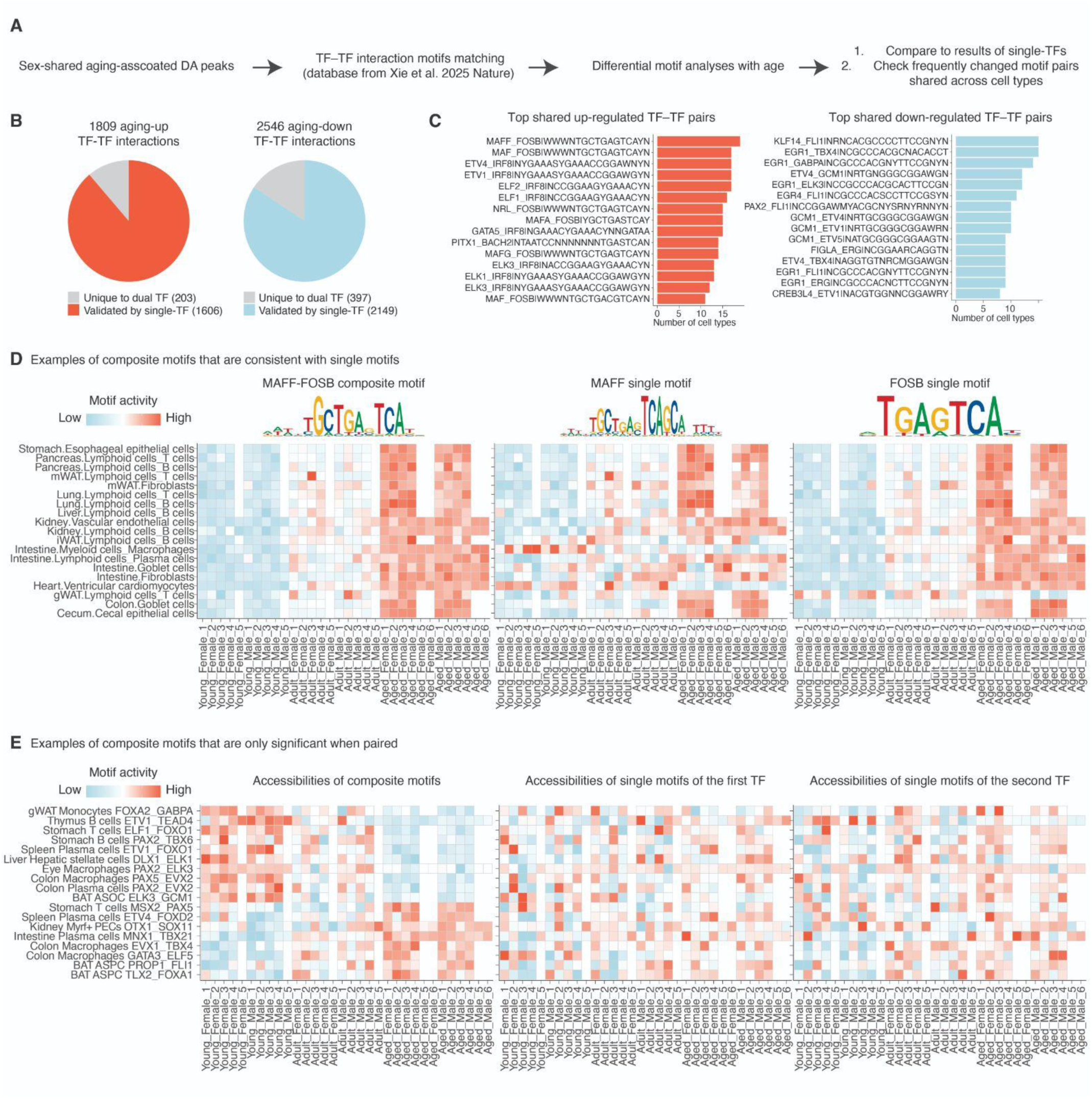
Combinatorial transcription factor interactions in aging-associated regulatory dynamics. (A) Analysis workflow. Sex-shared aging-associated DA peaks were intersected with TF–TF cooperative motifs identified from Xie. et al. Differential motif accessibility analyses were performed across ages for each cell type, followed by cross-cell-type summarization and comparison with single-TF results. **(B)** Pie charts showing the number of aging-up and aging-down TF–TF interactions, compared to validation by single TF motifs. TF-TF interactions were considered validated if either one of the pairs demonstrated significant changes with aging (FDR < 0.05) in the same direction. (C) Top recurrently altered TF–TF motif pairs across cell types, including up-regulated (left) and down-regulated (right) examples. (D) Heatmaps showing an example of a composite motif with consistent aging-associated accessibility changes across tissues. The increased accessibility pattern was also observed for the corresponding single-TF motifs. Each column represents one biological replicate. **(E)** Heatmaps illustrating composite motifs that are significant only when analyzed as pairs. Y-axis: motif names with corresponding cell types. X-axis: biological replicates. The first heatmap shows accessibility changes across ages for the composite motif, while the second and third show accessibility for each individual TF motif in the pair.

## Supplementary Tables

**Table S1.** Metadata for animals included in this study.

**Table S2.** List of gene markers used for cell type annotations.

**Table S3.** Differentially abundant main cell types with age across tissues. “qvalue_age” and “qvalue_interaction” indicate the significance of a non-zero coefficient for the age term and the age-sex interaction term respectively. “Regression R2” refers to the coefficient of determination of the regression model. “Pearson r” means the Pearson correlation r calculated between cell fraction of each main cell type within their corresponding tissues and ages, including samples of both sexes.

**Table S4.** Differentially abundant subtypes with age across tissues. “qvalue_age” and “qvalue_interaction” indicate the significance of a non-zero coefficient for the age term and the age-sex interaction term respectively. “Regression R2” refers to the coefficient of determination of the regression model. “Pearson r” means the Pearson correlation r calculated between cell fraction of each main cell type within their corresponding tissues and ages, including samples of both sexes.

**Table S5.** Differentially abundant immune subtypes with age across tissues. “qvalue_age” and “qvalue_interaction” indicate the significance of a non-zero coefficient for the age term and the age-sex interaction term respectively. “Regression R2” refers to the coefficient of determination of the regression model. “Pearson r” means the Pearson correlation r calculated between cell fraction of each main cell type within their corresponding tissues and ages, including samples of both sexes.

**Table S6.** Differentially abundant non-immune subtypes with age across tissues. “qvalue_age” and “qvalue_interaction” indicate the significance of a non-zero coefficient for the age term and the age-sex interaction term respectively. “Regression R2” refers to the coefficient of determination of the regression model. “Pearson r” means the Pearson correlation r calculated between cell fraction of each main cell type within their corresponding tissues and ages, including samples of both sexes.

**Table S7.** Differential accessible peaks with age shared in both sexes, including logFC, p-value, and q-value for each peak in both females and males.

**Table S8.** Differential accessible peaks with age unique to females, logFC, p-value, and q-value for each peak in both females and males.

**Table S9.** Differential accessible peaks with age unique to males, logFC, p-value, and q-value for each peak in both females and males.

**Table S10.** Aging-associated linkages with consistent changes in gene expression and chromatin accessibility at promoters and putative cis-regulatory elements. Each row represents a gene-peak-CRE linkage, including the gene name and genomic coordinates of the promoter and linked cis-regulatory site. The column “group” indicates whether the linkage is identified from both sexes or from one sex.

